# Effects of noise and metabolic cost on cortical task representations

**DOI:** 10.1101/2023.07.11.548492

**Authors:** Jake P. Stroud, Michał Wójcik, Kristopher T. Jensen, Makoto Kusunoki, Mikiko Kadohisa, Mark J. Buckley, John Duncan, Mark G. Stokes, Máté Lengyel

## Abstract

Cognitive flexibility requires both the encoding of task-relevant and the ignoring of task-irrelevant stimuli. While the neural coding of task-relevant stimuli is increasingly well understood, the mechanisms for ignoring task-irrelevant stimuli remain poorly understood. Here, we study how task performance and biological constraints jointly determine the coding of relevant and irrelevant stimuli in neural circuits. Using mathematical analyses and task-optimized recurrent neural networks, we show that neural circuits can exhibit a range of representational geometries depending on the strength of neural noise and metabolic cost. By comparing these results with recordings from primate prefrontal cortex (PFC) over the course of learning, we show that neural activity in PFC changes in line with a minimal representational strategy. Specifically, our analyses reveal that the suppression of dynamically irrelevant stimuli is achieved by activity-silent, sub-threshold dynamics. Our results provide a normative explanation as to why PFC implements an adaptive, minimal representational strategy.

## Introduction

How systems solve complex cognitive tasks is a fundamental question in neuroscience and artificial intelligence ^2–6^. A key aspect of complex tasks is that they often involve multiple types of stimuli, some of which can even be *irrelevant* for performing the correct behavioural response ^7–9^ or predicting reward ^10,11^. Over the course of task exposure, subjects must typically identify which stimuli are relevant and which are irrelevant. Examples of irrelevant stimuli include those that are irrelevant at all times in a task ^7,11–14^ — which we refer to as *static* irrelevance — and stimuli that are relevant at some time points but are irrelevant at other times in a trial — which we refer to as *dynamic* irrelevance (e.g. as is often the case for context-dependent decision-making tasks ^8,15–17^). Although tasks involving irrelevant stimuli have been widely used, it remains an open question as to how different types of irrelevant stimuli should be represented, in combination with relevant stimuli, to enable optimal task performance.

One may naively think that statically irrelevant stimuli should always be suppressed. However, stimuli that are currently irrelevant may be relevant in a future task. Furthermore, it is unclear whether dynamically irrelevant stimuli should be suppressed at all since the information is ultimately needed by the circuit. It may therefore be beneficial for a neural circuit to represent irrelevant information as long as no unnecessary costs are incurred and task performance remains high. Several factors could have a strong impact on whether irrelevant stimuli affect task performance. For example, levels of neural noise in the circuit as well as energy constraints and the metabolic costs of overall neural activity ^15,18–26^ can affect how stimuli are represented in a neural circuit. Indeed, both noise and metabolic costs are factors that biological circuits must contend with ^27–30^. Despite these considerations, models of neural population codes, including hand-crafted models and optimized artificial neural networks, typically use only a very limited range of the values of such factors ^4,8,19,26,31–35^ (but also see ^3,20^). Therefore, despite the success of recent comparisons between neural network models and experimental recordings ^4,8,19,22,26,31,34–36^, we may only be recovering very few out of a potentially large range of different representational strategies that neural networks could exhibit ^37^.

One challenge for distinguishing between different representational strategies, particularly when analysing experimental recordings, is that some stimuli may simply be represented more strongly than others. In particular, we might expect stimuli to be strongly represented in cortex *a priori* if they have previously been important to the animal. Indeed, being able to represent a given stimulus when learning a new task is likely a prerequisite for learning whether it is relevant or irrelevant in that particular context ^2,10^. Previously, it has been difficult to distinguish between whether a given representation existed *a priori* or emerged as a consequence of learning because neural activity is typically only recorded *after* a task has already been learned. A more relevant question is how the representation *changes* over learning ^11,38–41^, which provides insights into how the specific task of interest affects the representational strategy used by an animal or artificial network ^42^.

To resolve these questions, we optimized recurrent neural networks on a task that involved two types of irrelevant stimuli. One feature of the stimulus was statically irrelevant, and another feature of the stimulus was dynamically irrelevant. We found that, depending on the neural noise level and metabolic cost that was imposed on the networks during training, a range of representational strategies emerged in the optimized networks, from maximal (representing all stimuli) to minimal (representing only relevant stimuli). We then compared the strategies of our optimized networks with *learning-resolved* recordings from the prefrontal cortex (PFC) of monkeys exposed to the same task. We found that the representational geometry of the neural recordings changed in line with the minimal strategy. Using a simplified model, we derived mathematically how the strength of relevant and irrelevant coding depends on the noise level and metabolic cost. We then confirmed our theoretical predictions in both our task-optimized networks and neural recordings. By reverse-engineering our task-optimized networks, we also found that activity-silent, sub-threshold dynamics led to the suppression of dynamically irrelevant stimuli, and we confirmed predictions of this mechanism in our neural recordings.

In summary, we provide a mechanistic understanding of how different representational strategies can emerge in both biological and artificial neural circuits over the course of learning in response to salient biological factors such as noise and metabolic costs. These results in turn explain why PFC appears to employ a minimal representational strategy by filtering out task-irrelevant information.

## Results

### A task involving relevant and irrelevant stimuli

We study a task used in previous experimental work that uses a combination of multiple, relevant and irrelevant stimuli ^1^ (Fig. 1a). The task consists of an initial ‘fixation’ period, followed by a ‘color’ period, in which one of two colors are presented. After this, in the ‘shape’ period, either a square or diamond shape is presented (while the color stimulus stays on), such that the width of the shape can be either thick or thin. After this, the stimuli disappear and reward is delivered according to an XOR structure between color and shape (Fig. 1b). Note that the width of the shape is not predictive of reward, and it is therefore an *irrelevant* stimulus dimension (Fig. 1b). As width information is always irrelevant when it is shown, its irrelevance is *static*. In contrast, color is relevant during the shape period but could be ignored during the color period without loss of performance. Hence its irrelevance is *dynamic*.

**Fig. 1.**
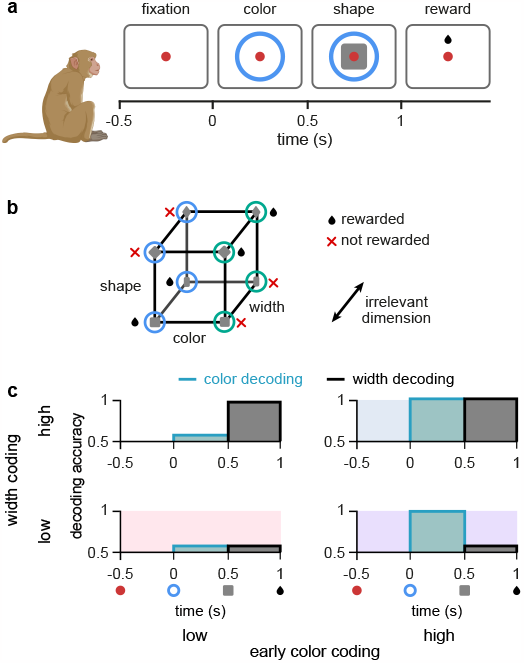
Task design and irrelevant stimulus representations. **a**, Illustration of the timeline of task events in a trial with the corresponding displays and names of task periods. Red dot shows fixation ring, blue (or green) circles appear during the color and shape periods, gray squares (or diamonds) appear during the shape period, and a juice reward is given during the reward period for rewarded combinations of color and shape stimuli (see panel b). No behavioural response was required for the monkeys as it was a passive object–association task ^1^. **b**, Schematic showing that rewarded conditions of color and shape stimuli follows an XOR structure. In addition, the width of the shape was not predictive of reward and was thus an irrelevant stimulus dimension. **c**, Schematic of 4 possible representational strategies, as indicated by linear decoding of population activity, for the task shown in panels a and b. Turquoise lines with shading show early color decoding and black lines with shading show width decoding. Strategies are split according to whether early color decoding is low (left column) or high (right column), and whether width decoding is low (bottom row) or high (top row).

Due to the existence of multiple different forms of irrelevant stimuli, there exist multiple different representational strategies for a neural circuit solving the task in Fig. 1a. These representational strategies can be characterised by assessing the extent to which different stimuli are linearly decodable from neural population activity ^10,14,43,44^. We use linear decodability because it only requires the computation of simple weighted sums of neural responses, and as such, it is a widely accepted criterion for the usefulness of a neural representation ^45^. Moreover, while representational strategies can differ along several dimensions in this task (for example, the decodability of color or shape during the shape period — both of which are task-relevant ^1^), our main focus here is on the two dimensions that specifically control the representation of irrelevant stimuli. Therefore, depending on whether each of the irrelevant stimuli are linearly decodable during their respective period of irrelevance, we distinguish four different (extreme) strategies, ranging from a ‘minimal strategy’, in which the irrelevant stimuli are only weakly represented (Fig. 1c, bottom left; pink shading), to a ‘maximal strategy’, in which both irrelevant stimuli are strongly represented (Fig. 1c, top right; blue shading).

### Stimulus representations in task-optimized recurrent neural networks

To understand the factors determining which representational strategy neural circuits employ to solve this task, we optimized recurrent neural networks to perform the task (Fig. 2a; Methods 1.2). The neural activities in these stochastic recurrent networks evolved according to

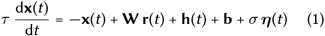

With

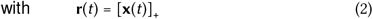

where **x**(*t*) = (*x*_1_(*t*), …, *x*_*N*_ (*t*))^*⊤*^ corresponds to the vector of ‘subthreshold’ activities of the *N* = 50 neurons in the network, **r**(*t*) is their momentary firing rates which is a rectified linear function (ReLU) of the subthreshold activity, *τ* = 50 ms is the effective time constant, and **h**(*t*) denotes the inputs to the network encoding the three stimulus features as they become available over the course of the trial (Fig. 2a, bottom). The optimized parameters of the network were **W**, the recurrent weight matrix describing connection strengths between neurons in the network (Fig. 2a, top; middle), **W**_in_, the feedforward weight matrix describing connections from the stimulus inputs to the network (Fig. 2a, top; left; see also Methods 1.2), and **b**, a stimulus-independent bias. Importantly, *σ* is the standard deviation of the neural noise process (Fig. 2a, top; pale gray arrows; with ***η***(*t*) being a sample from a Gaussian white noise process with mean 0 and variance 1), and as such represents a fundamental constraint on the operation of the network. The output of the network was given by

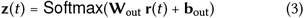

with optimized parameters **W**_out_, the readout weights (Fig. 2a, top; right), and **b**_out_, a readout bias.

**Fig. 2.**
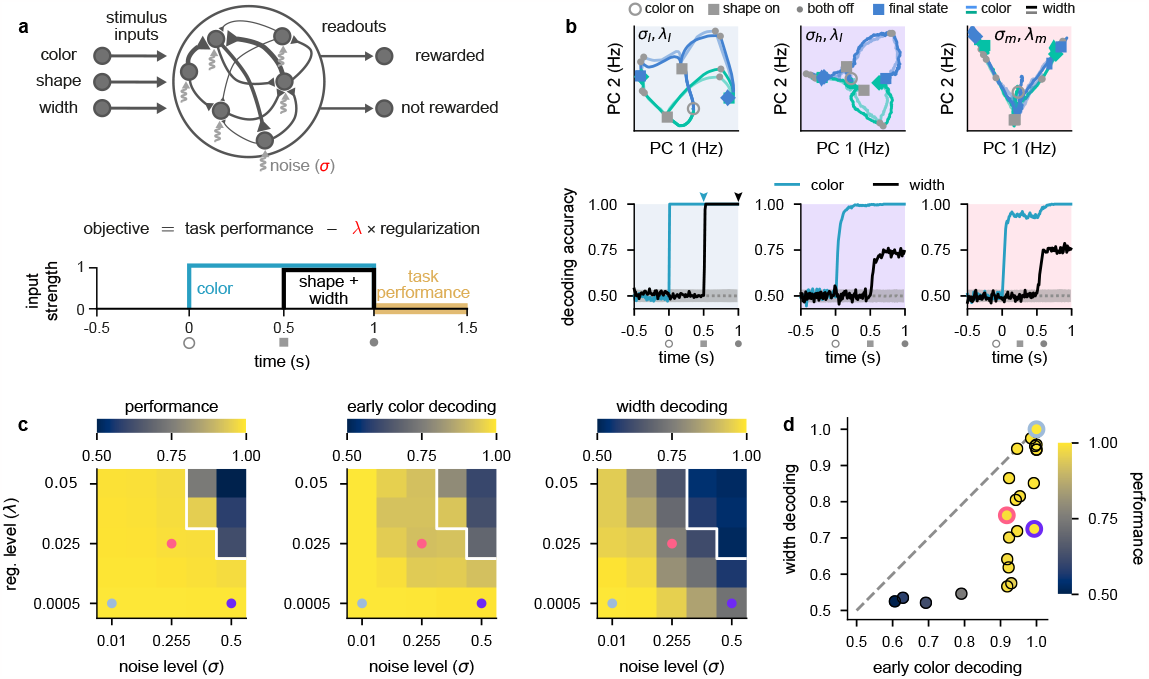
Stronger levels of noise and firing rate regularization lead to suppression of task-irrelevant stimuli in optimized recurrent networks. **a**, Top: illustration of a recurrent neural network model where each neuron receives independent white noise input with strength *σ* (middle). Color, shape, and width inputs are delivered to the network via 3 input channels (left). Firing rate activity is read out into two readout channels (either rewarded or not rewarded; right). All recurrent weights in the network, as well as weights associated with the input and readout channels, were optimized (Methods 1.2). Bottom: cost function used for training the recurrent neural networks (cf. Eq. 4) and timeline of task events within a trial for the recurrent neural networks (cf. Fig. 1a). Yellow shading on time axis shows the time period in which the task performance term enters the cost function (cf. Eq. 4). **b**, Top: neural firing rate trajectories in the top two PCs for an example network over the course of a trial (from color stimulus onset) for a particular noise (*σ*) and regularization (*λ*) regime. Open gray circles indicate color onset, filled gray squares indicate shape onset, filled gray circles indicate offset of both stimuli, and colored thick and thin squares and diamonds indicate the end of the trial at 1.5 s for all stimulus conditions. Pale and brightly colored trajectories indicate the two width conditions. We show results for networks exposed to low noise and low regularization (*σ*_l_, *λ*_l_; left, pale blue shading), high noise and low regularization (*σ*_h_, *λ*_l_; middle, purple shading), and medium noise and medium regularization (*σ*_m_, *λ*_m_; right, pink shading). Bottom: performance of a linear decoder (mean over 10 networks) trained at each time point within the trial to decode color (turquoise) or width (black) from neural firing rate activity for each noise and regularization regime. Dotted gray lines and shading show mean *±* 2 s.d. of chance level decoding based on shuffling trial labels. **c**, Left: performance of optimized networks determined as the mean performance over all trained networks during the reward period (Fig. 2a, bottom; yellow shading) for all noise (*σ*, horizontal axis) and regularization levels (*λ*, vertical axis) used during training. Pale blue, pink, and purple dots indicate parameter values that correspond to the dynamical regimes shown in panel b and Fig. 1c with the same background coloring. For parameter values above the white line, networks achieved a mean performance of less than 0.95. Middle: early color decoding determined as mean color decoding over all trained networks at the end of the color period (Fig. 2b, bottom left, turquoise arrow) using the same plotting scheme as the left panel. Right: width decoding determined as mean width decoding over all trained networks at the end of the shape period (Fig. 2b, bottom left, black arrow) using the same plotting scheme as the left panel. **d**, Width decoding plotted against early color decoding for all noise and regularization levels and colored according to performance. Pale blue, pink, and purple highlights indicate the parameter values shown with the same colors in panel c.

We optimized networks for a canonical cost function ^20,32,33,35,46^ (Fig. 2a, bottom; Methods 1.2.1):

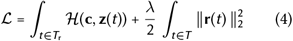

The first term in Eq. 4 is a task performance term measuring the cross-entropy loss ℋ (**c, z**(*t*)) between the one-hot encoded target choice, **c**, and the network’s output probabilities, **z**(*t*), during the reward period, *T*_r_ (Fig. 2a, bottom; yellow shading). The second term in Eq. 4 is a firing rate regularization term. This regularization term can be interpreted as a form of energy or metabolic cost ^18,24,31,46,47^ measured across the whole trial, *T*, because it penalizes large overall firing rates. Therefore, optimizing this cost function encourages networks to not only solve the task, but to do so using low overall firing rates. How important it is for the network to keep firing rates low is controlled by the ‘regularization’ parameter *λ*. Critically, we used different noise levels *σ* and regularization strengths *λ* (Fig. 2a, highlighted in red) to examine how these two constraints affected the dynamical strategies employed by the optimized networks to solve the task and compared them to our four hypothesised representational strategies (Fig. 1c). We focused on these two parameters because they are both critical factors for real neural circuits and *a priori* can be expected to have important effects on the resulting circuit dynamics. For example, metabolic costs will constrain the overall level of firing rates that can be used to solve the task while noise levels directly affect how reliably stimuli can be encoded in the network dynamics. For the remainder of this section, we analyse representational strategies utilized by networks after training. In subsequent sections, we analyse learning-related changes in both our optimized networks and neural recordings.

We found that networks trained in a low noise–low firing rate regularization setting (which we denote by *σ*_l_, *λ*_l_) employed a maximal representational strategy (Fig. 2b, left column; pale blue shading). Trajectories in neural firing rate space diverged for the two different colors as soon as the color stimulus was presented (Fig. 2b, top left; blue and green trajectories from open gray circle to gray squares), which resulted in high color decoding during the color period (Fig. 2b, bottom left; turquoise line; Methods 1.3.1). During the shape period, the trajectories corresponding to each color diverged again (Fig. 2b, top left; trajectories from gray squares to filled gray circles), such that all stimuli were highly decodable, including width (Fig. 2b, bottom left; black line, Supplementary Fig. 1a). After removal of the stimuli during the reward period, trajectories converged to one of two parts of state space according to the XOR task rule — which is required for high performance (Fig. 2b, top left; trajectories from filled gray circles to colored squares and diamonds). Because early color and width were highly decodable in these networks trained with a low noise and low firing rate regularization, the dynamical strategy they employed corresponds to the maximal representational regime (cf. Fig. 1c, blue shading).

We next considered the setting of networks that solve this task while being exposed to a high level of neural noise (which we denote by *σ*_h_, *λ*_l_; Fig. 2b, middle column; purple shading). In this setting, we also observed neural trajectories that diverged during the color period (Fig. 2b, top middle; gray squares are separated), which yielded high color decoding during the color period (Fig. 2b, bottom middle; turquoise line). However, in contrast to networks with a low level of neural noise (Fig. 2b, left), width was poorly decodable during the shape period (Fig. 2b, bottom middle; black line). Therefore, for networks challenged with higher levels of neural noise, the irrelevant stimulus dimension of width is represented more weakly (cf. Fig. 1c, purple shading). A similar representational strategy was also observed in networks that were exposed to a low level of noise but a high level of firing rate regularization (Supplementary Fig. 1c, black lines).

Finally, we considered the setting of networks that solve this task while being exposed to medium levels of both neural noise and firing rate regularization (which we denote by *σ*_m_, *λ*_m_; Fig. 2b, right column; pink shading). In this setting, neural trajectories diverged only weakly during the color period (Fig. 2b, top middle; gray squares on-top of one another), which yielded relatively poor (non-ceiling) color decoding during the color period (Fig. 2b, bottom right; turquoise line). Nevertheless, color decoding was still above chance despite the neural trajectories strongly overlapping in the 2-dimensional state space plot in Fig. 2b, top right), because these trajectories became separable in the full-dimensional state space of these networks. Additionally, width decoding was also poor during the shape period (Fig. 2b, bottom right; black line). Therefore, networks that were challenged with increased levels of both neural noise and firing rate regularization employed dynamics in line with a minimal representational strategy by only weakly representing irrelevant stimuli (cf. Fig. 1c, pink shading).

To gain a more comprehensive picture of the full range of dynamical solutions that networks can exhibit, we performed a grid search over multiple different levels of noise and firing rate regularization. Nearly all parameter values allowed networks to achieve high performance (Fig. 2c, left), except when both the noise and regularization levels were high (Fig. 2c, left; parameter values above white line). Importantly, all of the dynamical regimes that we showed in Fig. 2b achieved similarly high performances (Fig. 2c, left; pale blue, purple, and pink dots).

When looking at early color decoding (defined as color decoding at the end of the color period; Fig. 2b, bottom left, turquoise arrow) and width decoding (defined as width decoding at the end of the shape period; Fig. 2b, bottom left, black arrow), we saw a consistent pattern of results. Early color decoding was high only when either the lowest noise level was used (Fig. 2c, middle; *σ* = 0.01 column) or when the lowest regularization level was used (Fig. 2c, middle; *λ* = 0.0005 row). In contrast, width decoding was high only when both the level of noise and regularization were small (Fig. 2c, right; bottom left corner). Otherwise, width decoding became progressively worse as either the noise or regularization level was increased and achieved values that were typically lower than the corresponding early color decoding (Fig. 2c, compare right with middle). This pattern becomes clearer when we plot width decoding against early color decoding (Fig. 2d). No set of parameters yielded higher width decodability compared to early color decodability (Fig. 2d, no data point above the identity line). This means that we never observed the fourth dynamical regime we hypothesized *a priori*, in which width decoding would be high and early color decoding would be low (Fig. 1c, top left). Therefore, information with static irrelevance (width) was more strongly suppressed compared to information whose irrelevance was dynamic (color). We also note that we never observed pure chance levels of decoding of color or width during stimulus presentation. This is likely because it is challenging for recurrent neural networks to strongly suppress their inputs and it may also be the case that other hyperparameter regimes more naturally lead to stronger suppression of inputs (we discuss this second possibility later; e.g. Fig. 5).

### Comparing learning-related changes in stimulus representations in neural networks and primate lateral PFC

To understand the dynamical regime employed by PFC, we analyzed a data set ^1^ of multi-channel recordings from lateral prefrontal cortex (lPFC) in two monkeys exposed to the task shown in Fig. 1a. These recordings yielded 376 neurons in total across all recording sessions and both animals (Methods 1.1). Importantly, for understanding the direction in which neural geometries changed over learning, recordings commenced in the first session in which the animals were exposed to the task — that is, the recordings spanned the entirety of the learning process. For our analyses, we distinguished between the first half of recording sessions (Fig. 3a, gray; ‘early learning’) and the second half of recording sessions (Fig. 3a, black; ‘late learning’). A previous analysis of this dataset showed that, over the course of learning, the XOR representation of the task comes to dominate the dynamics during the late shape period of the task ^1^. Here however, we focus on relevant and irrelevant task variable coding during the stimulus periods and compare the recordings to the dynamics of task-optimized recurrent neural networks. Also, in line with the previous study ^1^, we do not examine the reward period of the task because the one-to-one relationship between XOR and reward in the data (which is not present in the models), will likely lead to trivial differences in XOR representations in the reward period between the data and models.

**Fig. 3.**
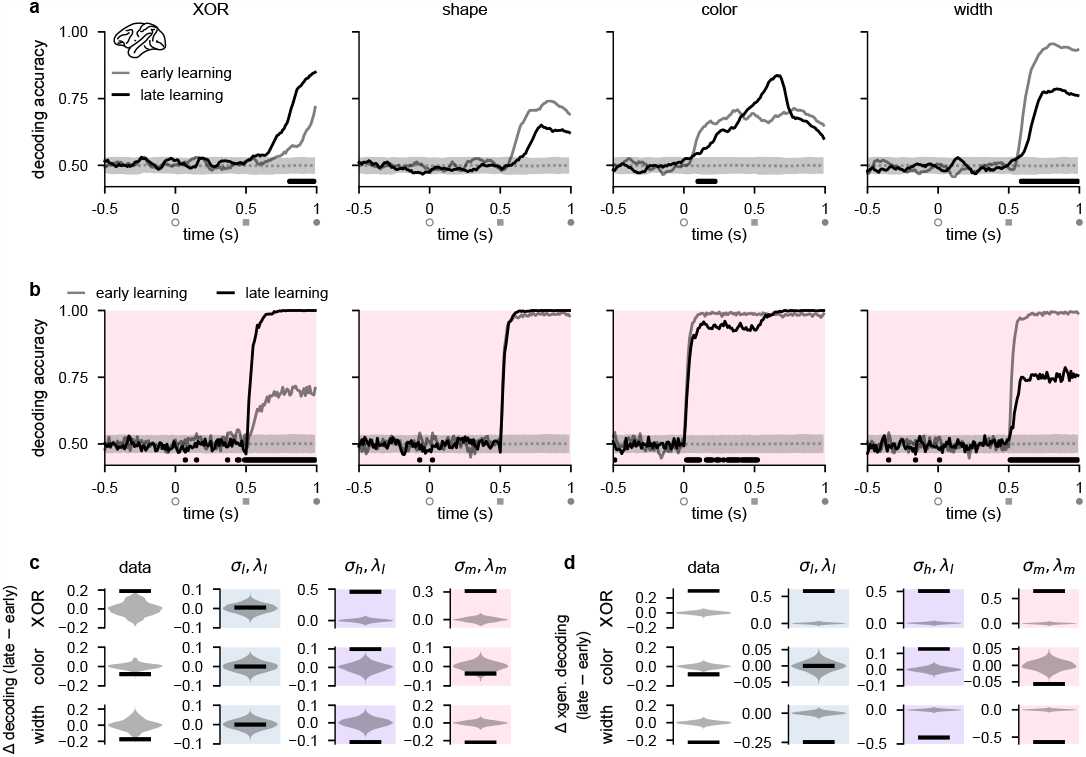
Stimulus representations in primate lateral PFC correspond to a minimal representational strategy. **a**, Performance of a linear decoder trained at each time point to predict either XOR (far left), shape (middle left), color (middle right), or width (far right) from neural population activity in lPFC (Methods 1.1 and 1.3.1). We show decoding results separately from the first half of sessions (gray, ‘early learning’) and the second half of sessions (black, ‘late learning’). Dotted gray lines and shading show mean *±* 2 s.d. of chance level decoding based on shuffling trial labels. Horizontal black bars show significant differences between early and late decoding using a two-sided cluster-based permutation test (Methods 1.4). Open gray circles, filled gray squares, and filled gray circles on the horizontal indicate color onset, shape onset, and offset of both stimuli, respectively. **b**, Same as panel a but for decoders trained on neural activity from optimized recurrent neural networks in the *σ*_m_, *λ*_m_ regime (cf. Fig. 2b–d, pink). **c**, Black horizontal lines show the mean change between early and late decoding during time periods when there were significant differences between early and late decoding in the data (horizontal black bars in panel a) for XOR (top row), color (middle row), and width (bottom row). (No significant differences in shape decoding were observed in the data; cf. panel a.) Violin plots show chance differences between early and late decoding based on shuffling trial labels. We show results for the data (far left column), *σ*_l_, *λ*_l_ networks (middle left column, pale blue shading), *σ*_h_, *λ*_l_ networks (middle right column, purple shading), and *σ*_m_, *λ*_m_ networks (far right column, pink shading). **d**, Same as panel c but we show results using cross-generalized decoding ^10^ during the same time periods as those used in panel c (Methods 1.3.1).

Similar to our analyses of our recurrent network models (Fig. 2b, bottom), we performed population decoding of the key task variables in the experimental data (Fig. 3a, Methods 1.3.1). We found that the decodability of the XOR relationship between task-relevant stimuli that determined task performance (Fig. 1b) significantly increased during the shape period over the course of learning, consistent with the animals becoming more familiar with the task structure (Fig. 3a, far left; compare gray and black lines). We also found that shape decodability during the shape period decreased slightly over learning (Fig. 3a, middle left; compare gray and black lines from gray square to gray circle), while color decodability during the shape period increased slightly (Fig. 3a, middle right; compare gray and black lines from gray square to gray circle). Importantly, however, color decodability significantly *decreased* during the color period (when it is ‘irrelevant’; Fig. 3a, middle right; compare gray and black lines), and width decodability significantly decreased during the shape period (Fig. 3a, far right; compare gray and black lines). Neural activities in lPFC thus appear to change in line with the minimal representational strategy over the course of learning, consistent with recurrent networks trained in the *σ*_m_, *λ*_m_ regime (cf. Fig. 1c, pink and Fig. 2b, bottom, pink).

We then directly compared these learning-related changes in stimulus decodability from lPFC with those that we observed in our task-optimized recurrent neural networks (cf. Fig. 2). We found that the temporal decoding dynamics of networks trained with medium noise and firing rate regularization (*σ*_m_, *λ*_m_; Fig. 2b– d, pink shading and pink dots) exhibited decoding dynamics most similar to those that we observed in lPFC. Specifically, XOR decodability significantly increased after shape onset, consistent with the networks learning the task (Fig. 3b, far left; compare gray and black lines). We also found that shape and color decodability did not significantly change during the shape period (Fig. 3b, middle left and middle right; compare gray and black lines from gray square to gray circle). Importantly, however, color decodability significantly *decreased* during the color period (when it is ‘irrelevant’; Fig. 3b, middle right; compare gray and black lines from open gray circle to gray square), and width decodability significantly decreased during the shape period (Fig. 3a, far right; compare gray and black lines). Other noise and regularization settings yielded temporal decoding that displayed a poorer resemblance to the data (Supplementary Fig. 1). For example, *σ*_l_, *λ*_l_ and *σ*_l_, *λ*_h_ networks exhibited almost no changes in decodability during the color and shape periods (Supplementary Fig. 1a,c) and *σ*_h_, *λ*_l_ networks exhibited increased XOR, shape, and color decodability at all times after stimulus onset while width decodability decreased during the shape period (Supplementary Fig. 1b). We also found that if regularization is driven to very high levels, color and shape decoding becomes weak during the shape period while XOR decoding remains high (Supplementary Fig. 1d). Therefore such networks effectively perform a pure XOR computation during the shape period.

We also note that the *σ*_m_, *λ*_m_ model does not perfectly match the decoding dynamics seen in the data. For example, although not significant, we observed a decrease in shape decoding and an increase in color decoding in the data during the shape period whereas the model only displayed a slight (non significant) increase in decodability of both shape and color during the same time period. These differences may be due to fundamentally different ways that brains encode sensory information upstream of PFC, compared to the more simplistic abstract sensory inputs used in models (see Discussion).

To systematically compare learning-related changes in the data and models, we analyzed time periods when there were significant changes in decoding in the data over the course of learning (Fig. 3a, horizontal black bars). This yielded a substantial increase in XOR decoding during the shape period, and substantial decreases in color and width decoding during the color and shape periods, respectively, in the data (Fig. 3c, far left column). During the same time periods in the models, networks in the *σ*_l_, *λ*_l_ regime exhibited no changes in decoding (Fig. 3c, middle left column; blue shading). In contrast, networks in the *σ*_h_, *λ*_l_ regime exhibited substantial increases in XOR and color decoding, and a substantial decrease in width decoding (Fig. 3c, middle right column; purple shading). Finally, in line with the data, networks in the *σ*_m_, *λ*_m_ regime exhibited a substantial increase in XOR decoding, and substantial decreases in both color and width decoding (Fig. 3c, middle right column; pink shading).

In addition to studying changes in traditional decoding, we also studied learning-related changes in ‘cross-generalized decoding’ which provides a measure of how factorized the representational geometry is across stimulus conditions ^10^ (Supplementary Fig. 2). (For example, for evaluating cross-generalized decoding for color, a color decoder trained on square trials would be tested on diamond trials, and vice versa ^10^; Methods 1.3.1.) Using this measure, we found that changes in decoding were generally more extreme over learning and that models and data bore a stronger resemblance to one another compared with traditional decoding. Specifically, all models and the data showed a strong increase in XOR decoding (Fig. 3d, top row, ‘XOR’) and a strong decrease in width decoding (Fig. 3d, bottom row, ‘width’). However, only the data and *σ*_m_, *λ*_m_ networks showed a decrease in color decoding (Fig. 3d, compare middle far left and middle far right), whereas *σ*_l_, *λ*_l_ networks showed no change in color decoding (Fig. 3d, middle left; blue shading) and *σ*_h_, *λ*_l_ networks showed an increase in color decoding (Fig. 3d, middle right; purple shading).

Beside studying the effects of input noise and firing rate regularization, we also examined the effects of different strengths of the initial stimulus input connections prior to training ^15^. In line with previous studies ^3,20,35,46,48^, for all of our previous results, the initial input weights were set to small random values prior to training (Methods 1.2.1). We found that changing these weights had similar effects to changing neural noise (with the opposite sign). Specifically, when input weights were set to 0 before training, initial decoding was at chance levels and only increased with learning for XOR, shape, and color (whereas width decoding hardly changed with learning and remained close to chance levels; Supplementary Fig. 3) — analogously to the *σ*_h_, *λ*_l_ regime (Supplementary Fig. 1b). In contrast, when input weights were set to large random values prior to training, initial decoding was at ceiling levels and did not significantly change over learning during the color and shape periods for any of the task variables (Supplementary Fig. 4) — similar to what we found in the *σ*_l_, *λ*_l_ regime (Supplementary Fig. 1a). Thus, neither extremely small nor extremely large initial input weights were consistent with the data that exhibited both increases and decreases in decodability of task variables over learning (Fig. 3a).

### Theoretical predictions for strength of relevant and irrelevant stimulus coding

To gain a theoretical understanding of how irrelevant stimuli should be processed in a neural circuit, we performed a mathematical analysis of a minimal linear model that only included a single relevant stimulus and a single statically irrelevant stimulus (Supplement A1). Although this analysis applies to a simpler task compared with that faced by our neural networks and animals, crucially it still allows us to understand how relevant and irrelevant coding depend on noise and metabolic constraints. Our mathematical analysis suggested that the effects of noise and firing rate regularization on the performance and metabolic cost of a network can be understood via three key aspects of its representation: the strength of neural responses to the relevant stimulus (‘relevant coding’), the strength of responses to the irrelevant stimulus (‘irrelevant coding’), and the overlap between the population responses to relevant and irrelevant stimuli (‘overlap’; Fig. 4a). In particular, maximizing task performance (i.e. the decodability of the relevant stimulus) required relevant coding to be strong, irrelevant coding to be weak, and the overlap between the two to be small (Fig. 4b, top; Supplement A1.3). This ensures that the irrelevant stimulus interferes minimally with the relevant stimulus. In contrast, to reduce a metabolic cost (such as we considered previously in our optimized recurrent networks, see Eq. 4 and Fig. 2), both relevant and irrelevant coding should be weak (Fig. 4b, bottom; Supplement A1.4).

**Fig. 4.**
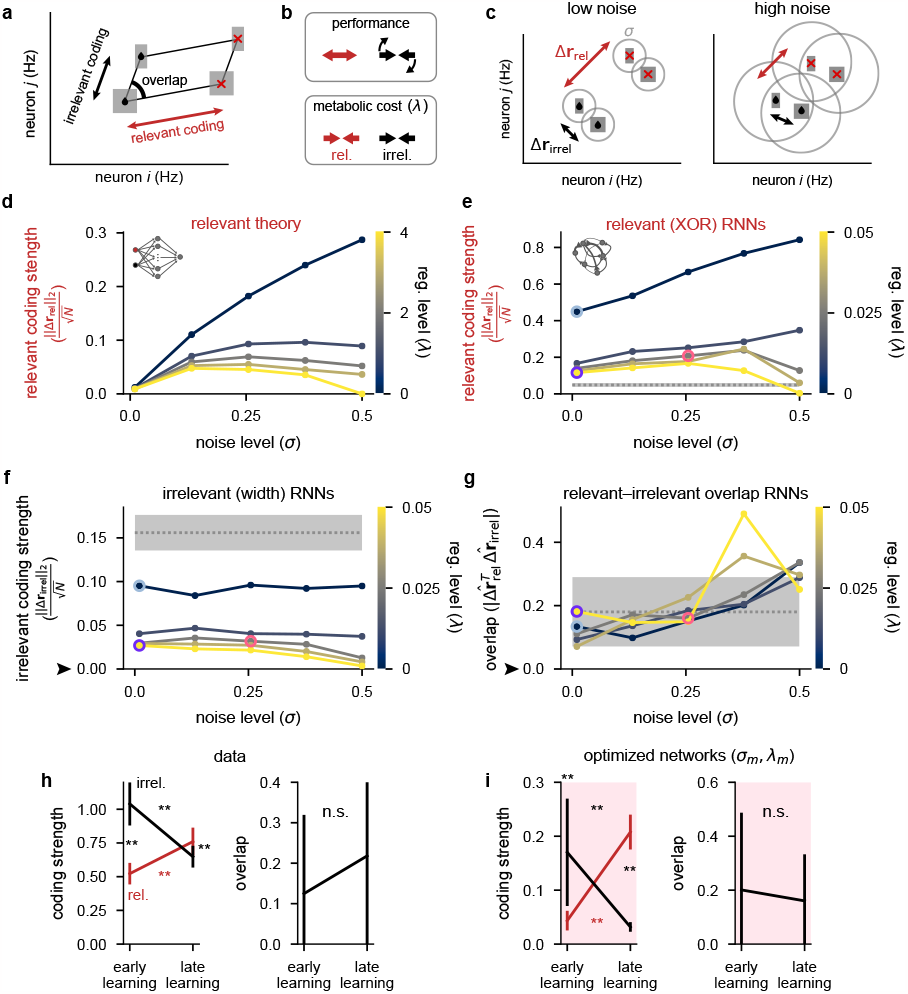
Theoretical predictions for strength of relevant and irrelevant stimulus coding and comparison to lPFC recordings. **a**, Schematic of activity in neural state space for two neurons for a task involving two relevant (black drops vs. red crosses) and two irrelevant stimuli (thick vs. thin squares). Strengths of relevant and irrelevant stimulus coding are shown with red and black arrows, respectively, and the overlap between relevant and irrelevant coding is also shown (‘overlap’). **b**, Schematic of our theoretical predictions for the optimal setting of relevant (red arrows) and irrelevant (black arrows) coding strengths when either maximizing performance (top) or minimizing a metabolic cost with strength *λ* (bottom; see Eq. 4). **c**, Schematic of our theoretical predictions for the strength of relevant (Δ**r**_rel_; black drops vs. red crosses; red arrows) and irrelevant (Δ**r**_irrel_; thick vs. thin squares; black arrows) coding when jointly optimizing for both performance and a metabolic cost (cf. panel b). In the low noise regime (left), relevant conditions are highly distinguishable and irrelevant conditions are poorly distinguishable as well as strongly orthogonal to the relevant conditions. In the high noise regime (right), all conditions are poorly distinguishable. **d**, Our theoretical predictions (Eq. S34) for the strength of relevant coding (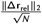, see panel c) as a function of the noise level *σ* (horizontal axis) and firing rate regularization strength *λ* (colorbar). **e**, Same as panel d but for our optimized recurrent neural networks ( Fig. 2) where we show the strength of relevant (XOR) coding (Methods 1.3.2). Pale blue, purple, and pink highlights correspond to the noise and regularization strengths shown in Fig. 2c,d. Gray dotted line and shading shows mean*±*2 s.d. (over 250 networks; 10 networks for each of the 25 different noise and regularization levels) of 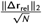 prior to training. **f**, Same as panel e but for the strength of irrelevant (width) coding 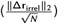. The black arrow indicates the theoretical prediction of 0 irrelevant coding. **g**, The absolute value of the normalized dot product (overlap) between relevant and irrelevant representations 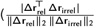, i.e. 0 implies perfect orthogonality and 1 implies perfect overlap) for our optimized recurrent neural networks. The black arrow indicates the theoretical prediction of 0 overlap. **h**, Left: coding strength (length of arrows in panel a; Methods 1.3.2) for relevant (XOR; red) and irrelevant (width; black) stimuli during early and late learning for our lPFC recordings. Right: the absolute value of the overlap between relevant and irrelevant representations for our lPFC recordings (0 implies perfect orthogonality and 1 implies perfect overlap). Error bars show the mean *±* 2 s.d. over 10 non-overlapping splits of the data. **i**, Same as panel c but for the optimized recurrent neural networks in the *σ*_m_, *λ*_m_ regime (see pink dots in Fig. 2). Error bars show the mean *±* 2 s.d. over 10 different networks. P-values resulted from a two-sided Mann–Whitney U test (*, *p <* 0.05; **, *p <* 0.01; n.s., not significant; see Methods 1.4).

In combination, when decoding performance and metabolic cost are *jointly* optimized, as in our taskoptimized recurrent networks (Fig. 2), our theoretical analyses suggested that performance should decrease with both the noise level *σ* and the strength of firing rate regularization *λ* in an approximately interchangeable way, and metabolic cost should increase with *σ* but decrease with *λ* (Supplement A1.5). We also found that the strength of relevant coding should decrease with *λ*, but its dependence on *σ* was more nuanced. For small *σ*, the performance term effectively dominates the metabolic cost (Fig. 4b, top) and the strength of relevant coding should increase with *σ*. However, if *σ* is too high, the strength of relevant coding starts decreasing otherwise a disproportionately high metabolic cost must be incurred to achieve high performance (Fig. 4c,d). Our mathematical analysis also suggested that irrelevant coding and relevant– irrelevant overlap could in principle depend on the noise and metabolic cost strength — particularly if performing noisy optimization where the curvature of the cost function can be relatively shallow around the minimum (Supplement A1.5). Therefore, practically, we also expect that irrelevant coding should be mostly dependent (inversely) on *λ*, but relevant–irrelevant overlap should mostly depend on *σ* (Supplement A1.6). These theoretical predictions were confirmed by our recurrent neural network simulations (Fig. 4e–g).

We next measured, in both recorded and simulated population responses, the three aspects of population responses that our theory identified as being key in determining the performance and metabolic cost of a network (Fig. 4a; Methods 1.3.2). We found a close correspondence in the learning-related changes of these measures between our lPFC recordings (Fig. 4h) and optimized recurrent networks (Fig. 4i). In particular, we found that the strength of relevant (XOR) coding increased significantly over the course of learning (Fig. 4h,i, left; red). The strength of irrelevant (width) coding decreased significantly over the course of learning (Fig. 4h,i, left; black), such that it became significantly smaller than the strength of relevant coding (Fig. 4h,i, left; compare red and black at ‘late learning’). Finally, relevant and irrelevant directions were always strongly orthogonal in neural state space, and the level of orthogonality did not significantly change with learning (Fig. 4h,i, right). Although we observed no learning-related changes in overlap, it may be that for stimuli that are more similar than the relevant and irrelevant features we studied here (XOR and width), the overlap between these features may decrease over learning rather than simply remaining small.

### Activity-silent, sub-threshold dynamics lead to the suppression of dynamically irrelevant stimuli

While the representation of statically irrelevant stimuli can be suppressed by simply weakening the input connections conveying information about it to the network, the suppression of dynamically irrelevant stimuli requires a mechanism that alters the dynamics of the network (since this information ultimately needs to be used by the network to achieve high performance). To gain some intuition about this mechanism, we first analyzed 2-neuron networks trained on the task. To demonstrate the basic mechanism of suppression of dynamically irrelevant stimuli (i.e. weak early color coding), we compared the *σ*_l_, *λ*_l_ (Fig. 5a) and *σ*_m_, *λ*_m_ (Fig. 5b) regimes for these networks, as these corresponded to the minimal and maximal amount of suppression of dynamically irrelevant stimuli (Fig. 2c).

**Fig. 5.**
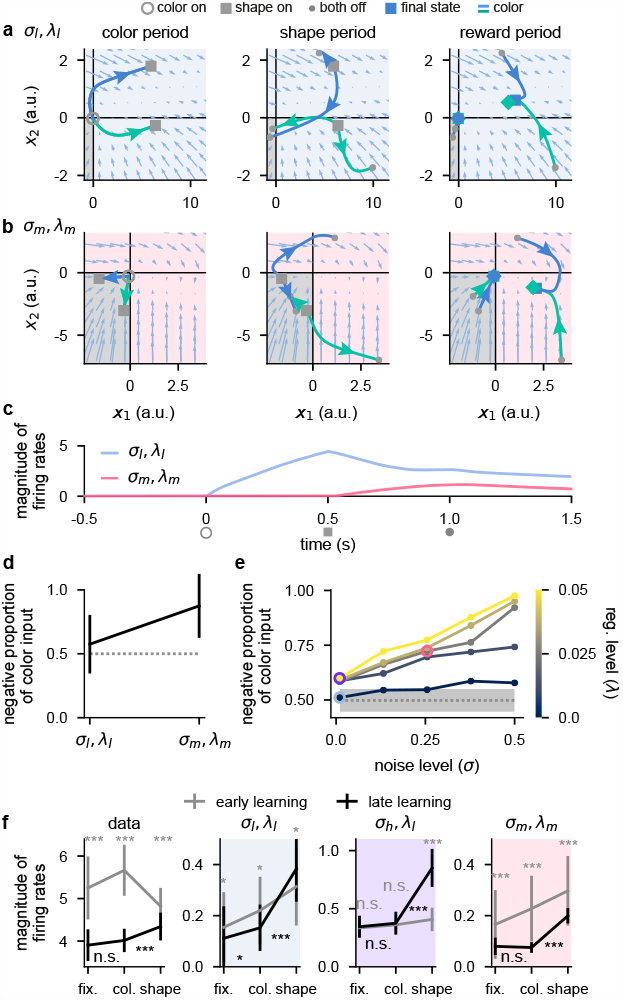
Activity-silent, sub-threshold dynamics lead to the suppression of dynamically irrelevant stimuli. **a**, Sub-threshold neural activity (**x**(*t*) in Eq. 1) in the full state space of an example 2-neuron network over the course of a trial (from color onset) trained in the *σ*_l_, *λ*_l_ regime. Pale blue arrows show flow field dynamics (direction and magnitude of movement in the state space as a function of the momentary state). Open gray circles indicate color onset, gray squares indicate shape onset, filled gray circles indicate offset of both stimuli, and colored squares and diamonds indicate the end of the trial at 1.5 s. We plot activity separately for the 3 periods of the task (color period, left; shape period, middle; reward period, right). We plot dynamics without noise for visual clarity. **b**, Same as panel a but for a network trained in the *σ*_m_, *λ*_m_ regime. **c**, Momentary magnitude of firing rates 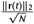 i.e. momentary metabolic cost, see Eq. 4 ) for the 2-neuron networks from panels a (blue line) and b (pink line). **d**, Mean *±* 2 s.d. (over 10 2-neuron networks) proportion of color inputs that have a negative sign for the two noise and regularization regimes shown in panels a–c. Gray dotted line shows chance level proportion of negative color input. **e**, Mean (over 10 50-neuron networks) proportion of color inputs that have a negative sign for all noise (horizontal axis) and regularization (colorbar) strengths shown in Fig. 2c,d. Pale blue, purple, and pink highlights correspond to the noise and regularization strengths shown in Fig. 2c,d. Gray dotted line and shading shows mean±2 s.d. (over 250 networks; 10 networks for each of the 25 different noise and regularization levels) of the proportion of negative color input prior to training (i.e. the proportion of negative color input expected when inputs are drawn randomly from a Gaussian distribution; Methods 1.2.1). **f**, Momentary magnitude of firing rates 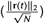 for our lPFC recordings (far left) and 50-neuron networks in the *σ*_l_, *λ*_l_ regime (middle left, blue shading), *σ*_h_, *λ*_l_ regime (middle right, purple shading), and *σ*_m_, *λ*_m_ regime (far right, pink shading). Error bars show the mean *±* 2 s.d. over either 10 non-overlapping data splits for the data or over 10 different networks for the models. P-values resulted from a two-sided Mann–Whitney U test (*, *p <* 0.05; **, *p <* 0.01; ***, *p <* 0.001; n.s., not significant; see Methods 1.4).

We examined trajectories of sub-threshold neural activity **x**(*t*) (cf. Eq. 1) in the full 2-neuron state space (Fig. 5a,b, blue and green curves). We distinguished between the negative quadrant of state space, which corresponds to the rectified part of the firing rate nonlinearity (Fig. 5a,b, bottom left gray quadrant; cf. Eq. 2), and the rest of state space. Importantly, if sub-threshold activity lies within the negative quadrant at some time point, both neurons in the network have zero firing rate and consequently a decoder cannot decode any information from the firing rate activity and the network exhibits no metabolic cost (cf. Eq. 4). Therefore, we reasoned that color inputs may drive sub-threshold activity so that it lies purely in the negative quadrant of state space so that no metabolic cost is incurred during the color period (akin to nonlinear gating ^15,49^). Later in the trial, when these color inputs are combined with the shape inputs, activity may then leave the negative quadrant of state space so that the network can perform the task.

We found that for the *σ*_l_, *λ*_l_ regime, there was typically at least one set of stimulus conditions for which subthreshold neural activity evolved outside the negative quadrant of state space during any task period (Fig. 5a). Consequently, color was decodable to a relatively high level during the color period (Supplementary Fig. 5a, middle right) and this network produced a relatively high metabolic cost throughout the task (Fig. 5c, blue line). In contrast, for networks in the *σ*_m_, *λ*_m_ regime, the two color inputs typically drove neural activity into the negative quadrant of state space during the color period (Fig. 5b, left). Therefore, during the color period, the network produced zero firing rate activity (Fig. 5c, pink line from open gray circle to gray square). Consequently, color was poorly decodable (Supplementary Fig. 5b, middle right; black line) and the network incurred no metabolic cost during the color period. Thus, color information was represented in a sub-threshold, ‘activity-silent’ ^50^ state during the color period. However, during the shape and reward periods later in the trial, the color inputs, now in combination with the shape inputs, affected the firing rate dynamics and the neural trajectories explored the full state space in a similar manner to the *σ*_l_, *λ*_l_ network (Fig. 5b, middle and right panels). Indeed, we also found that color decodability increased substantially during the shape period in the *σ*_m_, *λ*_m_ network (Supplementary Fig. 5b, middle right; black line). This demonstrates how color inputs can cause no change in firing rates during the color period when they are irrelevant, and yet these same inputs can be utilized later in the trial to enable high task performance (Supplementary Fig. 5a,b, far left). While we considered individual example networks in Fig. 5a–c, color inputs consistently drove neural activity into the negative quadrant of state space across repeated training of *σ*_m_, *λ*_m_ networks but not for *σ*_l_, *λ*_l_ networks (Fig. 5d).

Next, we performed the same analysis as Fig. 5d on the large (50-neuron) networks that we studied previously (Figs. 2–4). Similar to the 2-neuron networks, we found that large networks in the *σ*_l_, *λ*_l_ regime did not exhibit more negative color inputs than would be expected by chance (Fig. 5e, pale blue highlighted point) — consistent with the high early color decoding we found previously in these networks (Fig. 2c, middle, pale blue highlighted point). However, when increasing either the noise or regularization level, the proportion of negative color inputs increased above chance such that for the highest noise and regularization level, nearly all color inputs were negative (Fig. 5e). We also found a strong negative correlation between the proportion of negative color input and the level of early color decoding that we found previously in Fig. 2c (Supplementary Fig. 5c, *r* = − 0.9, *p <* 10^−9^). This suggests that color inputs that drive neural activity into the rectified part of the firing rate nonlinearity, and thus generate purely sub-threshold activity-silent dynamics, is the mechanism that generates weak early color coding during the color period in these networks. We also examined whether these results generalized to networks that use a sigmoid (as opposed to a ReLU) nonlinearity. To do this, we trained networks with a tanh nonlinearity (shifted for a meaningful comparison with ReLU, so that the lower bound on firing rates was 0, rather than− 1) and found qualitatively similar results to the ReLU networks. In particular, color inputs drove neural activity towards 0 firing rate during the color period in *σ*_m_, *λ*_m_ networks but not in *σ*_l_, *λ*_l_ networks (Supplementary Fig. 6, compare a and b), which resulted in a lower metabolic cost during the color period for *σ*_m_, *λ*_m_ networks compared to *σ*_l_, *λ*_l_ networks (Supplementary Fig. 6c). This was reflected in color inputs being more strongly negative in *σ*_m_, *λ*_m_ networks compared to *σ*_l_, *λ*_l_ networks which only showed chance levels of negative color inputs (Supplementary Fig. 6d).

We next sought to test whether this mechanism could also explain the decrease in color decodability over learning that we observed in the lPFC data. To do this, we measured the magnitude of firing rates in the fixation, color, and shape periods for both early and late learning (note that the magnitude of firing rates coincides with our definition of the metabolic cost; cf. Eq. 4 and Fig. 5c). To decrease metabolic cost over learning, we would expect two changes: firing rates should decrease with learning and firing rates should not significantly increase from the fixation to the color period after learning (Fig. 5c, pink line). Indeed, we found that firing rates decreased significantly over the course of learning in all task periods (Fig. 5f, far left; compare gray and black lines), and this decrease was most substantial during the fixation and color periods (Fig. 5f, far left). We also found that after learning, firing rates during the color period were not significantly higher than during the fixation period (Fig. 5f, far left; compare black error bars during fixation and color periods). During the shape period however, firing rates increased significantly compared to those during the fixation period (Fig. 5f, far left; compare black error bars during fixation and shape periods). Therefore, the late learning dynamics of the data are in line with what we saw for the optimized 2-neuron *σ*_m_, *λ*_m_ network (Fig. 5c, pink line).

We then compared these results from our neural recordings with the results from our large networks trained in different noise and regularization regimes. We found that for *σ*_l_, *λ*_l_ networks, over the course of learning, firing rates decreased slightly during the fixation and color periods but actually increased during the shape period (Fig. 5f, middle left pale blue shading; compare gray and black lines). Additionally, after learning, firing rates increased between fixation and color periods and between color and shape periods (Fig. 5f, middle left pale blue shading; compare black error bars). For *σ*_h_, *λ*_l_ networks, over the course of learning, we found that firing rates did not significantly change during the fixation and color periods but increased significantly during the shape period (Fig. 5f, middle right purple shading; compare gray and black lines). Furthermore, after learning, firing rates did not change significantly between fixation and color periods but did increase significantly between color and shape periods (Fig. 5f, middle right purple shading; compare black error bars). Finally, for the *σ*_m_, *λ*_m_ networks, we found a pattern of results that was most consistent with the data. Firing rates decreased over the course of learning in all task periods (Fig. 5f, far right pink shading; compare gray and black lines) and the decrease in firing rates was most substantial during the fixation and color periods. After learning, firing rates did not change significantly from fixation to color periods but did increase significantly during the shape period (Fig. 5f, far right; compare black error bars).

To investigate whether these findings depended on our random network initialization prior to training, we also compared late learning firing rates to firing rates that resulted from randomly shuffling the color, shape, and width inputs (which emulates alternative tasks where different combinations of color, shape, and width are relevant). For example, the relative strengths of the 3 inputs to the network prior to training on this task may affect how the firing rates change over learning. Under this control, we also found that *σ*_*m*_, *λ*_*m*_ networks bore a close resemblance to the data (Supplementary Fig. 5d, compare black lines to blue lines).

## Discussion

Comparing the neural representations of task-optimized networks with those observed in experimental data has been particularly fruitful in recent years ^4,8,19,26,31,34–36^. However, networks are typically optimized using only a very limited range of hyperparameter values ^4,8,19,26,31–35^. Instead, here we showed that different settings of key, biologically relevant hyperparameters such as noise and metabolic costs, can yield a variety of qualitatively different dynamical regimes that bear varying degrees of similarity with experimental data. In general, we found that increasing levels of noise and firing rate regularization led to increasing amounts of irrelevant information being filtered out in the networks. Indeed, filtering out of task-irrelevant information is a well-known property of the PFC and has been observed in a variety of tasks ^7,12,31,38,49,51–53^. We provide a mechanistic understanding of the specific conditions that lead to stronger filtering of task-irrelevant information. We predict that these results should also generalize to richer, more complex cognitive tasks that may, for example, require context switching ^15,38,52^ or invoke working memory ^7,13,52^. Indeed, filtering out of task-irrelevant information in the PFC has been observed in such tasks ^7,13,15,38,52,54^.

Our results are also likely a more general finding of neural circuits that extend beyond the PFC. In line with this, it has previously been shown that strongly regularized neural network models trained to reproduce monkey muscle activities during reaching bore a stronger resemblance to neural recordings from primary motor cortex compared to unregularized models ^19^. In related work on motor control, recurrent networks controlled by an optimal feedback controller recapitulated key aspects of experimental recordings from primary motor cortex (such as orthogonality between preparatory and movement neural activities) when the control input was regularized ^24^. Additionally, regularization of neural firing rates, and its natural biological interpretation as a metabolic cost, has recently been shown to be a key ingredient for the formation of grid cell-like response profiles in artificial networks ^18,22^.

By showing that PFC representations changed in line with a minimal representational strategy, our results are in line with various studies showing lowdimensional representations under a variety of tasks in the PFC and other brain regions ^13,15,31,55,56^. This is in contrast to several previous observations of highdimensional neural activity in PFC ^2,10^. Both high- and low-dimensional regimes confer distinct yet useful benefits: high-dimensional representations allow many behavioural readouts to be generated, thereby enabling highly flexible behaviour ^2,15,26,57–59^, whereas low-dimensional representations are more robust to noise and allow for generalization across different stimuli ^15,26,59^. These two different representational strategies have previously been studied in models by setting the initial network weights to either small values (to generate low-dimensional ‘rich’ representations) or large values ^15^ (to generate high-dimensional ‘lazy’ representations). However, in contrast to previous approaches, we studied the more biologically plausible effects of firing rate regularization (i.e. a metabolic cost; see also the supplement of Ref. 15) on the network activities over the course of learning and compared them to learning-related changes in PFC neural activities. Firing rate regularization will cause neural activities to wax and wane as networks are exposed to new tasks depending upon the stimuli that are currently relevant. In line with this, it is conceivable that prolonged exposure to a task that has a limited number of stimulus conditions, some of which can even be generalized over (as was the case for our task), encourages more low dimensional dynamics to form ^1,15,48,59–61^. In contrast, tasks that use a rich variety of stimuli (that may even dynamically change over the task ^4,6,62^), and which do not involve generalization across stimulus conditions, may more naturally lead to high-dimensional representations ^2,26,48,59,61,63^. It would therefore be an important future question to understand how our results also depend on the task being studied as some tasks may more naturally lead to the ‘maximal’ representational regime ^26,48,61^ (Figs. 1– 3 and 5, blue shading).

A previous analysis of the same dataset that we studied here focused on the late parts of the trial ^1^. In particular, they found that the final result of the computation needed for the task, the XOR operation between color and shape, emerges and eventually comes to dominate lPFC representations over the course of learning in the late shape period ^1^. Our analysis goes beyond this by studying mechanisms of suppression of both static and dynamically irrelevant stimuli across all task periods and how different levels of neural noise and metabolic cost can lead to qualitatively different representations of irrelevant stimuli in task-optimized recurrent networks. Other previous studies focused on characterizing the representation of several task-relevant ^2,10,14^ (and, in some cases, -irrelevant ^15^) variables over the course of individual trials. Characterizing how key aspects of neural representations change over the course of learning, as we did here, offers unique opportunities for studying the functional objectives shaping neural representations ^42^.

There were several aspects of the data that were not well captured by our models. For example, during the shape period, decodability of shape decreased while decodability of color increased (although not significantly) in our neural recordings (Fig. 3a). These differences in changes in decoding may be due to fundamentally different ways that brains encode sensory information upstream of PFC, compared to our models. For example, shape and width are both geometric features of the stimulus, whose encoding is differentiated from that of color already at early stages of visual processing ^64^. Such a hierarchical representation of inputs may automatically lead to the (un)learning about the relevance of width (which the *σ*_m_, *λ*_m_ model reproduced) generalizing to shape, but not to color. In contrast, inputs in our model used a non-hierarchical one-hot encoding (Fig. 2), which did not allow for such generalization. Moreover, in the data, we may expect width to be *a priori* more strongly represented than color or shape because it is a much more potent sensory feature. In line with this, in our neural recordings, we found that width was very strongly represented in early learning compared to the other stimulus features (Fig. 3a, far right) and width always yielded high cross-generalized decoding — even after learning (Supplementary Fig. 2a, far right). Nevertheless, studying *changes* in decoding over learning, rather than absolute decoding levels, allowed us to focus on features of learning that do not depend on the details of upstream sensory representations of stimuli. Future studies could incorporate aspects of sensory representations that we ignored here by using stimulus inputs with which the model more faithfully reproduces the experimentally observed initial decodability of stimulus features.

In line with previous studies ^3,18–20,24,31–33,35,46,47^, we operationalized metabolic cost in our models through *L*_2_ firing rate regularization. This cost penalizes high overall firing rates. There are however alternative conceivable ways to operationalize a metabolic cost; for example *L*_1_ firing rate regularization has been used previously when optimizing neural networks and promotes more sparse neural firing ^3^. Interestingly, although our *L*_2_ is generally conceived to be weaker than *L*_1_ regularization, we still found that it encouraged the network to use purely sub-threshold activity in our task. The regularization of synaptic weights may also be biologically relevant ^3^ because synaptic transmission uses the most energy in the brain compared to other processes ^65,66^. Additionally, even subthreshold activity could be regularized as it also consumes energy (although orders of magnitude less than spiking ^67^). Therefore, future work will be needed to examine how different metabolic costs affect the dynamics of task-optimized networks.

We build on several previous studies that have also analysed learning-related changes in PFC activity ^1,38,39,63^ — although these studies typically used reversal-learning paradigms in which animals are already highly task proficient and the effects of learning and task switching are inseparable. For example, in a rule-based categorization task in which the categorization rule changed after learning an initial rule, neurons in mouse PFC adjusted their selectivity depending on the rule such that currently irrelevant information was not represented ^38^. Similarly, neurons in rat PFC transition rapidly from representing a familiar rule to representing a completely novel rule through trial and error learning ^39^. Additionally, the dimensionality of PFC representations was found to increase as monkeys learned the value of novel stimuli ^63^. Importantly, however, PFC representations did not distinguish between novel stimuli when they were first chosen. It was only once the value of the stimuli were learned that their representations in PFC were distinguishable ^63^. These results are consistent with our results where we found poor XOR decoding during the early stages of learning which then increased over learning as the monkeys learned the rewarded conditions (Fig. 3a, far left). However, we also observed high decoding of width during early learning which was not predictive of reward (Fig. 3a, far right). One key distinction between our study and that of Ref. 63, is that our recordings commenced from the first trial the monkeys were exposed to the task. In contrast, in Ref. 63, the monkeys were already highly proficient at the task and so the neural representation of their task was already likely strongly task specific by the time recordings were taken.

In line with our approach here, several recent studies have also examined the effects of different hyperparameter settings on the solution that optimized networks exhibit. One study found that decreasing regularization on network weights led to more sequential dynamics in networks optimized to perform working memory tasks ^20^. Another study found that the number of functional clusters that a network exhibits does not depend strongly on the strength of (*L*_1_ rate or weight) regularization, but did depend upon whether the single neuron nonlinearity saturates at high firing rates ^3^. It has also been shown that networks optimized to perform path integration can exhibit a range of different properties, from grid cell-like receptive fields to distinctly non grid cell-like receptive fields, depending upon biologically relevant hyperparameters — including noise and regularization ^18,22,37^. Indeed, in addition to noise and regularization, various other hyperparameters have also been shown to affect the representational strategy used by a circuit, such as the firing rate nonlinearity ^3,18,37^ and network weight initialization ^15,37^. It is therefore becoming increasingly clear that analysing the interplay between learning and biological constraints will be key for understanding the computations that various brain regions perform.

## Acknowledgements

This work was funded by the Wellcome Trust (Investigator Award in Science 212262/Z/18/Z to M.L., Sir Henry Wellcome Postdoctoral Fellowship 215909/Z/19/Z to J.S., and award 101092/Z/13/Z to M.K., M.K., and J.D.), the Human Frontier Science Programme (Research Grant RGP0044/2018 to M.L.), the Medical Research Council UK Program (MC_UU_00030/7 to M.K., M.K., and J.D.), the Biotechnology and Biological Sciences Research Council (award BB/M010732/1 to M.G.S.), a Gates Cambridge scholarship (to K.T.J.), a Clarenden scholarship and Saven European scholarship (to M.W.), and the James S. McDonnell Foundation (award 220020405 to M.G.S.). For the purpose of open access, the authors have applied a CC-BY public copyright licence to any author accepted manuscript version arising from this submission. We thank Christopher Summerfield for useful feedback and detailed comments on the manuscript.

## Author Contributions

J.P.S., M.W., J.D., and M.G.S. conceived the study. J.D. and M.J.B. led the experimental recordings and M.W., M.K., and M.K. performed the experimental recordings and data pre-processing. J.P.S., K.T.J, and M.L. developed the theoretical framework and performed analytical derivations. J.P.S. performed all numerical simulations, and produced the figures. J.P.S. and M.W. performed the data analysis. J.P.S., M.W., K.T.J., and M.L. interpreted the results. All authors revised the final manuscript.

## Code availability

All code was custom written in Python using NumPy, SciPy, Matplotlib, Scikit-learn, and Tensorflow libraries. Code is available in the following repository: https://github.com/jakepstroud.

## 1 Methods

### 1.1 Experimental materials and methods

Experimental methods have been described previously ^1^. Briefly, two adult male rhesus macaques (monkey 1 and monkey 2) performed a passive object–association task (Fig. 1a,b; see the main text ‘A task involving relevant and irrelevant stimuli’ for a description of the task). Neural recordings commenced from the first session the animals were exposed to the task. All trials with fixation errors were excluded. The dataset contained on average 237.9 (s.d. = 23.9) and 104.8 (s.d. = 2.3) trials for each of the 8 conditions for monkeys 1 and 2, respectively. Data were recorded from the ventral and dorsal lateral PFC over a total of 27 daily sessions across both monkeys which yielded 146 and 230 neurons for monkey 1 and monkey 2, respectively. To compute neural firing rates, we convolved binary spike trains with a Gaussian kernel with a standard deviation of 50 ms. In order to characterize changes in neural dynamics over learning, analyses were performed separately on the first half of sessions (‘early learning’; 9 and 5 sessions from monkey 1 and monkey 2, respectively) and the second half of sessions (‘late learning’; 8 and 5 sessions from monkey 1 and monkey 2, respectively; Figs. 3–5 and Supplementary Fig. 2a).

### 1.2 Neural network models

The dynamics of our simulated networks evolved according to Eqs. 1 and 2 and are repeated here:

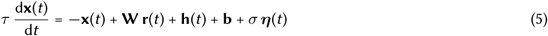

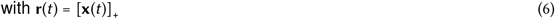

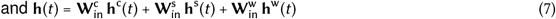

where **x**(*t*) = (*x*_1_(*t*), …, *x*_*N*_ (*t*))^*⊤*^ corresponds to the vector of ‘subthreshold’ activities of the *N* neurons in the network, **r**(*t*) is their momentary firing rates, obtained as a rectified linear function (ReLU) of their subthreshold activities (Eq. 6; except for the networks of Supplementary Fig. 6 in which we used a tanh nonlinearity to examine the generalizability of our results), *τ* = 50 ms is the effective time constant, **W** is the recurrent weight matrix, **h**(*t*) denotes the total stimulus input, **b** is a stimulus-independent bias, *σ* is the standard deviation of the neural noise process, ***η***(*t*) is a sample from a Gaussian white noise process with mean 0 and variance 1, 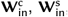 and 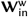 are color, shape, and width input weights, respectively, and **h**^c^(*t*), **h**^s^(*t*), and **h**^w^(*t*) are one-hot encodings of color, shape, and width inputs, respectively.

All simulations started at *t*_0_ = − 0.5 s and lasted until *t*_max_ = 1.5 s, and consisted of a fixation (− 0.5–0 s), color (0–0.5 s), shape (0.5–1 s), and reward (1–1.5 s) period (Fig. 1a and Fig. 2a). The initial condition of neural activity was set to **x**(*t*_0_) = **0**. In line with the task, elements of **h**^c^(*t*) were set to 0 outside the color and shape periods, and elements of both **h**^s^(*t*) and **h**^w^(*t*) were set to 0 outside the shape period. All networks used *N* = 50 neurons (except for Fig. 5a–d and Supplementary Fig. 5a,b which used *N* = 2 neurons). We solved the dynamics of Eqs. 5-7 using a first-order Euler–Maruyama approximation with a discretization time step of 1 ms.

#### 1.2.1 Network optimization

Choice probabilities were computed through a linear readout of network activities:

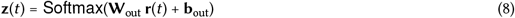

where **W**_out_ are the readout weights and **b**_out_ is a readout bias. To measure network performance, we used a canonical cost function ^3,20,32,33,35,46^ (Eqs. 3 and 4). We repeat the cost function from the main text here:

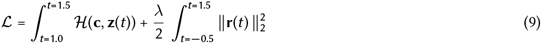

where the first term is a task performance term which consists of the cross-entropy loss 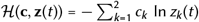 between the one-hot encoded target choice, **c** (based on the stimuli of the given trial, as defined by the task rules, Fig. 1b), and the network’s readout probabilities, **z**(*t*) (Eq. 8). Note that we measure total classification performance (cross entropy loss) during the reward period (integral in the first term runs from *t* = 1.0 to *t* = 1.5; Fig. 2a, bottom; yellow shading), as appropriate for the task. The second term in Eq. 9 is a widely-used ^3,18–20,24,32,33,35,46^ *L*_2_ regularization term (with strength *λ*) applied to the neural firing rates throughout the trial (integral in the second term runs from *t* = −0.5 to *t* = 1.5).

We initialized the free parameters of the network (the elements of 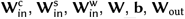, and 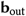) by sampling (independently) from a normal distribution of mean 0 and standard deviation 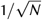 . There were two exceptions to this: we also investigated the effects of initializing the elements of the input weights 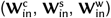 to either 0

(Supplementary Fig. 3) or sampling their elements from a normal distribution of mean 0 and standard deviation 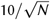 (Supplementary Fig. 4). After initialization, we optimized these parameters using gradient descent with Adam ^68^, where gradients were obtained from backpropagation through time. We used a learning rate of 0.001 and trained networks for 1000 iterations using a batch size of 10. For each noise *σ* and regularization level *λ* (see Table 1), we optimized 10 networks with different random initializations of the network parameters.

**Table 1.** Parameters used in the simulations of our models.

| Symbol | Fig. 2, Fig. 3b–d, Fig. 4e–g,i, Fig. 5e,f, Supplementary Figs. 1, 3 and 4, Supplementary Fig. 2b,c, and Supplementary Fig. 5c,d | Fig. 5a–d and Supplementary Fig. 5a,b | Supplementary Fig. 6 | Supplementary Fig. 1d | Units | Description |
| --- | --- | --- | --- | --- | --- | --- |
| $N$ | 50 | 2 | 2 | 50 | - | number of neurons |
| $t_0$ | -0.5 | -0.5 | -0.5 | -0.5 | s | simulation start time |
| $t_{\max}$ | 1.5 | 1.5 | 1.5 | 1.5 | s | simulation end time |
| $\tau$ | 0.05 | 0.05 | 0.05 | 0.05 | s | effective time constant |
| $r(\mathbf{x}(t))$ | ReLU | ReLU | $\tanh(\mathbf{x}(t)) + 1$ | ReLU | Hz | nonlinearity |
| $\sigma$ | variable<sup>a</sup> | [0.01, 0.255] | [0.01, 0.255] | 0.01 | - | noise standard deviation |
| $\lambda$ | variable<sup>b</sup> | [0.0005, 0.02525] | [0.0005, 0.02525] | 0.5 | s | regularization strength |
| $\mathbf{W}$ | optimized<sup>c</sup> | optimized<sup>c</sup> | optimized<sup>c</sup> | optimized<sup>c</sup> | s | weight matrix |
| $\mathbf{W}_{\text{in}}^c$ | optimized<sup>c</sup> | optimized<sup>c</sup> | optimized<sup>c</sup> | optimized<sup>c</sup> | - | color input weights |
| $\mathbf{W}_{\text{in}}^s$ | optimized<sup>c</sup> | optimized<sup>c</sup> | optimized<sup>c</sup> | optimized<sup>c</sup> | - | shape input weights |
| $\mathbf{W}_{\text{in}}^w$ | optimized<sup>c</sup> | optimized<sup>c</sup> | optimized<sup>c</sup> | optimized<sup>c</sup> | - | width input weights |
| $\mathbf{b}$ | optimized<sup>c</sup> | optimized<sup>c</sup> | optimized<sup>c</sup> | optimized<sup>c</sup> | - | bias |
| $\mathbf{W}_{\text{out}}$ | optimized<sup>c</sup> | optimized<sup>c</sup> | optimized<sup>c</sup> | optimized<sup>c</sup> | s | readout weights |
| $\mathbf{b}_{\text{out}}$ | optimized<sup>c</sup> | optimized<sup>c</sup> | optimized<sup>c</sup> | optimized<sup>c</sup> | - | readout bias |

<sup>a</sup> The noise standard deviation $\sigma$ took one of 5 evenly spaced values between 0.01 and 0.5 (see Fig. 2c).
<sup>b</sup> The firing rate regularization strength $\lambda$ took one of 5 evenly spaced values between 0.0005 and 0.05 (see Fig. 2c).
<sup>c</sup> See Methods 1.2 for details.

### 1.3 Analysis methods

Here we describe methods that we used to analyse neural activities. Whenever applicable, the same processing and analysis steps were applied to both experimentally recorded and model simulated data. All neural firing rates were sub-sampled at a 10-ms resolution and, unless stated otherwise, we did not trial-average firing rates before performing analyses. Analyses were either performed at every time point in the trial (Fig. 2b, Fig. 3a,b, Fig. 5a–c, Supplementary Figs. 1–4, and Supplementary Fig. 5a,b), at the end of either the color (Fig. 2c,d, and Supplementary Fig. 5c; ‘early color decoding’) or shape periods (Fig. 2c,d, ‘width decoding’), during time periods of significant changes in decoding over learning in the data (Fig. 3c,d), during the final 100 ms of the shape period (Fig. 4e–i), or during the final 100 ms of the fixation, color, and shape periods (Fig. 5f, and Supplementary Fig. 5d).

#### 1.3.1 Linear decoding

For decoding analyses (Fig. 2b–d, Fig. 3, Supplementary Figs. 1–4, and Supplementary Fig. 5a–c), we fitted decoders using linear support vector machines to decode the main task variables: color, shape, width, and the XOR between color and shape. We measured decoding performance in a cross-validated way, using separate sets of trials to train and test the decoders, and we show results averaged over 10 random 1:1 train:test splits. For firing rates resulting from simulated neural networks, we used 10 trials for both the train and test splits. Chance level decoding was always 0.5 as all stimuli were binary.

For showing changes in decoding over learning (Fig. 3c,d), we identified time periods of significant changes in decoding during the color and shape periods in the data (Fig. 3a, horizontal black bars; see Methods 1.4), and show the mean change in decoding during these time periods for both the data and models (Fig. 3c, horizontal black lines). We used the same time periods when showing changes in cross-generalized decoding over learning (Fig. 3d, see below) For cross-generalized decoding ^10^ (Fig. 3d and Supplementary Fig. 2), we used the same decoding approach as described above, except that cross validation was performed across trials corresponding to different stimulus conditions. Specifically, following the approach outlined in Ref. 10, because there were 3 binary stimuli in our task (color, shape, and width), there were 8 different trial conditions (Fig. 1b). Therefore, for each task variable (we focused on color, shape, width, and the XOR between color and shape), there were 6 different ways of choosing 2 of the 4 conditions corresponding to each of the two possible stimuli for that task variable (e.g. color 1 vs. color 2). For example, when training a decoder to decode color, there were 6 different ways of choosing 2 conditions corresponding to color 1, and 6 different ways of choosing 2 conditions corresponding to color 2 (the remaining 4 conditions were then used for testing the decoder). Therefore, for each task variable, there were 6 *×* 6 = 36 different ways of creating training and testing sets that corresponded to different stimulus conditions. We then took the mean decoding accuracy we obtained across all 36 different training and testing sets.

#### 1.3.2 Measuring stimulus coding strength

In line with our mathematical analysis (Supplement A1, and in particular Eq. S1) and in line with previous studies ^8,48,69^, to measure the strength of stimulus coding for relevant and irrelevant stimuli, we fitted the following linear regression model

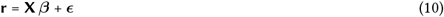

where **r** is size *K × N* (where *K* is the number of trials) and corresponds to the neural firing rates, **X** is size *K×* 3 where the first column is all 1 (thus encoding the mean firing rate of each neuron across all conditions) and elements of the final two columns are either − 0.5 or 0.5 depending upon whether trials correspond to XOR 1 or XOR 2 (relevant) conditions (column 2) or whether trials correspond to width 1 or width 2 (irrelevant) conditions (column 3). The coefficients to be fitted (***β***) is size 3 *× N* and has the following structure ***β*** = [***μ***, Δ**r**_rel_, Δ**r**_irrel_]^*⊤*^ where ***μ*** is the mean across all conditions, Δ**r**_rel_ are the coefficients corresponding to relevant (XOR) conditions, and Δ**r**_irrel_ are the coefficients corresponding to irrelevant (width) conditions. Finally, ***ϵ*** is size *K× N* and contains the residuals. Note that calculating the mean difference in firing rate between the two relevant conditions and between the two irrelevant conditions would yield identical estimations of Δ**r**_rel_ and Δ**r**_irrel_ because our stimuli are binary. (We also fitted decoders to decode either relevant or irrelevant conditions and extracted their coefficients to instead obtain Δ**r**_rel_ and Δ**r**_irrel_ and obtained near-identical results.)

We then calculated the Euclidean norm of both Δ**r**_rel_ and Δ**r**_irrel_ normalized by the number of neurons 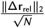 and 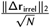 and the absolute value of the normalized dot product (overlap) between them 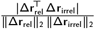 Fig. 4e–i ). For our neural recordings (Fig. 4h), we calculated these quantities separately for 10 non-overlapping splits of the data. For our neural networks (Fig. 4e–g,i, ), we calculated these quantities separately for 10 different networks.

#### 1.3.3 Measuring the magnitude of neural firing rates

We first calculated the firing rate for each condition and time point and then calculated the mean (across conditions and time points) Euclidean norm of firing rates appropriately normalized by the number of neurons: 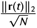 (Fig. 5f, and Supplementary Fig. 5d). For our neural recordings (Fig. 5f, far left), we calculated this separately for 10 non-overlapping splits of the data. For our neural networks (Fig. 5f, middle left, middle right, and far right, and Supplementary Fig. 5d), we calculated this separately for 10 different networks. For our optimized neural networks, to emulate alternative tasks where different combinations of color, shape, and width are relevant, we also performed the same analysis when shuffling all 6 input weights (after learning) in all 14 possible ways (Supplementary Fig. 5d, blue lines, ‘shuffle inputs’). These 14 possible shuffles excluded the original set up of input weights, any re-ordering of inputs within a single input channel (since any re-ordering within a single input channel would be identical up to a re-labelling), and any re-ordering between the shape and width input channels (since any re-ordering within the shape and width input channels would also be identical up to a re-labelling).

### 1.4 Statistics

For decoding analyses, we used non-parametric permutation tests to calculate statistical significance. We used 100 different random shuffles of condition labels across trials to generate a null distribution for decoding accuracy. We plotted chance level decoding (Fig. 3a,b, Supplementary Figs. 1–4, and Supplementary Fig. 5a,b) by combining both early and late learning null distributions.

To calculate significant differences in decoding accuracy over learning (Fig. 3a,b, Supplementary Figs. 1–4, and Supplementary Fig. 5a,b), our test statistic was the difference in decoding accuracy between early and late learning, and our null distribution was the difference in decoding accuracy between early and late learning for the 100 different shuffles of condition labels (see above). We calculated two-tailed p-values for all tests. Additionally, to control for time-related cluster-based errors, we also added a cluster-based permutation correction ^70^.

For all other tests, we used a two-sided Mann–Whitney U test (Fig. 4h,i, Fig. 5f, and Supplementary Fig. 5d).

## Supplementary figurse

**Supplementary Fig. 1.**
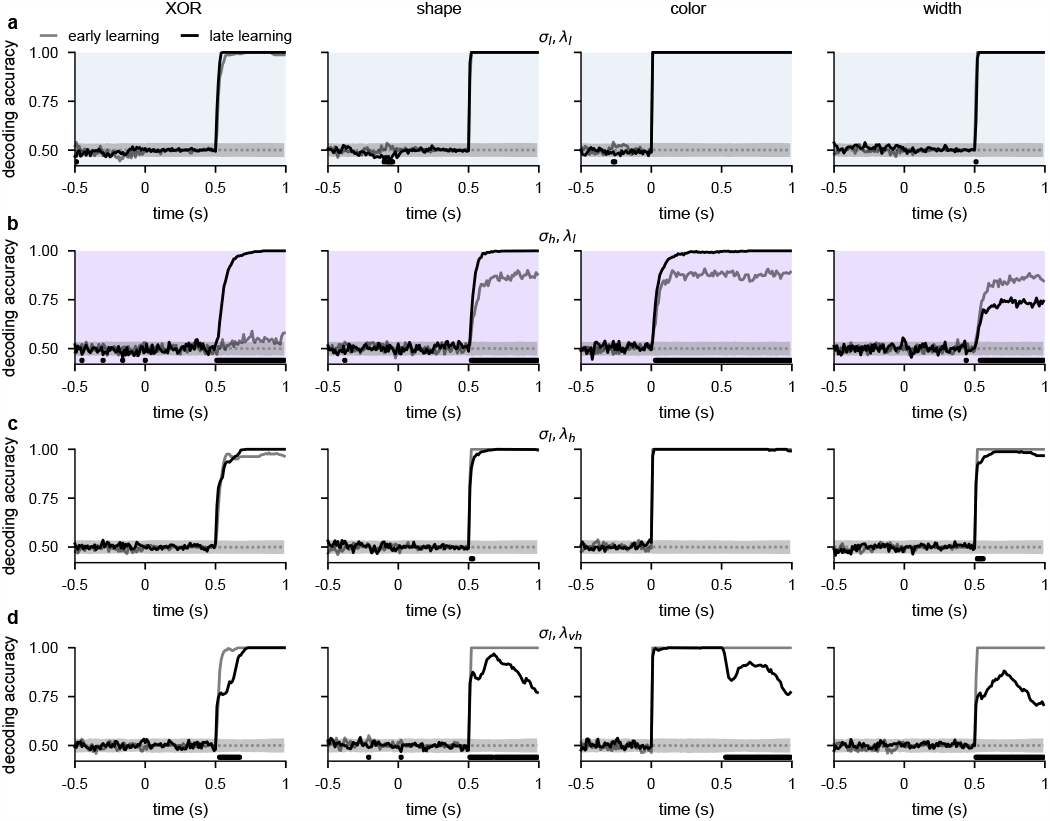
Temporal decoding for networks with different noise and regularization levels. **a**, Performance of a linear decoder trained at each time point to predict either XOR (far left), shape (middle left), color (middle right), or width (far right) from neural population activity from optimized recurrent neural networks trained in the *σ*_l_, *λ*_l_ regime. We show decoding results separately from the first half of sessions (gray, ‘early learning’) and the second half of sessions (black, ‘late learning’). Dotted gray lines and shading show mean *±* 2 s.d. of chance level decoding based on shuffling trial labels. Horizontal black bars show significant differences between early and late decoding using a two-sided cluster-based permutation test (Methods 1.4). **b**, Same as panel a, but for networks optimized in the *σ*_h_, *λ*_l_ regime. **c**, Same as panel a, but for networks optimized in the *σ*_l_, *λ*_h_ regime. **d**, Same as panel a, but for networks optimized in a very high regularization regime (*σ* = 0.01, *λ* = 0.5) while still being able to perform the task to a high accuracy.

**Supplementary Fig. 2.**
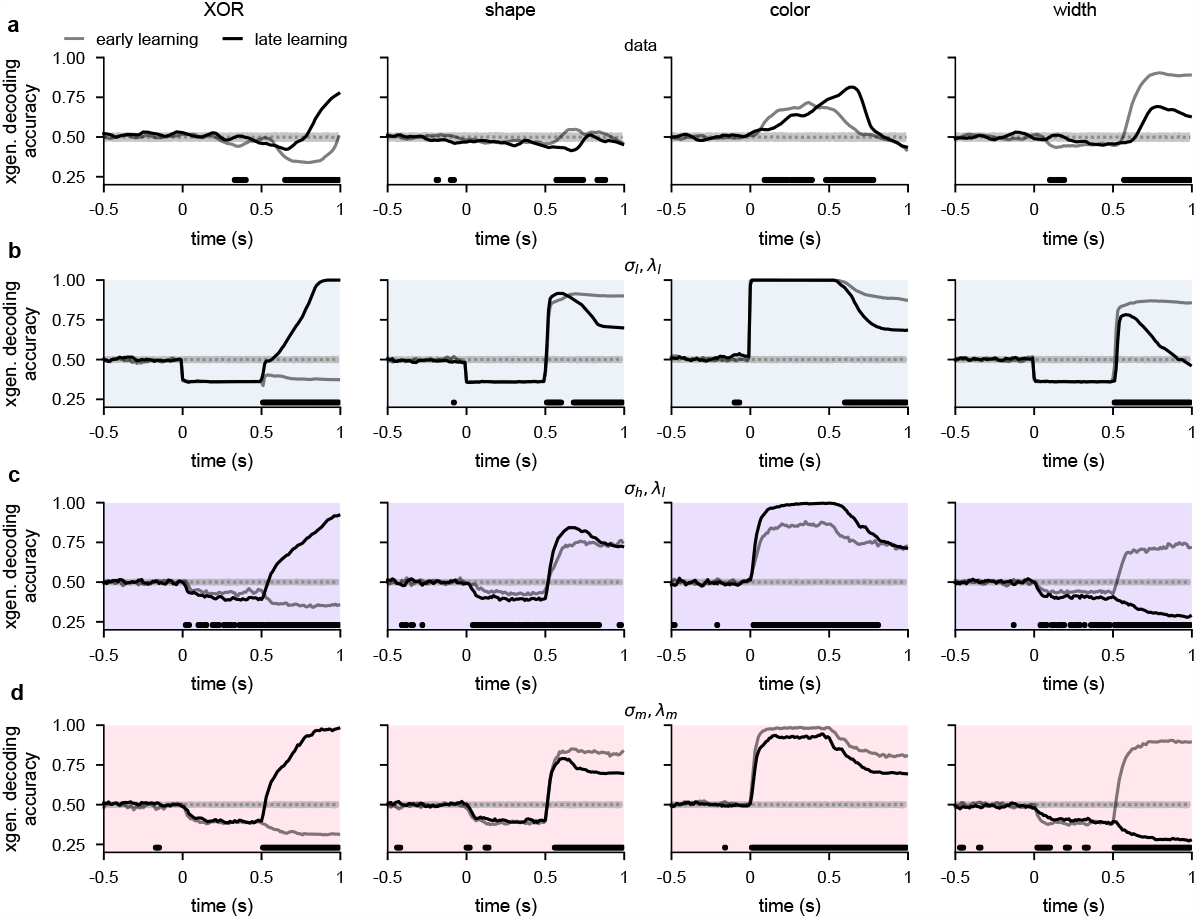
Cross-generalized temporal decoding for lPFC recordings and networks with different noise and regularization levels. **a**, Performance of a cross-generalized decoder (Methods 1.3.1) trained at each time point to predict either XOR (far left), shape (middle left), color (middle right), or width (far right) from neural population activity from our lPFC recordings. We show decoding results separately from the first half of sessions (gray, ‘early learning’) and the second half of sessions (black, ‘late learning’). Dotted gray lines and shading show mean *±* 2 s.d. of chance level decoding based on shuffling trial labels. Horizontal black bars show significant differences between early and late decoding using a two-sided cluster-based permutation test (Methods 1.4). **b**, Same as panel a, but for recurrent neural networks optimized in the *σ*_l_, *λ*_l_ regime. **c**, Same as panel a, but for recurrent neural networks optimized in the *σ*_h_, *λ*_l_ regime. **d**, Same as panel a, but for recurrent neural networks optimized in the *σ*_m_, *λ*_m_ regime.

**Supplementary Fig. 3.**
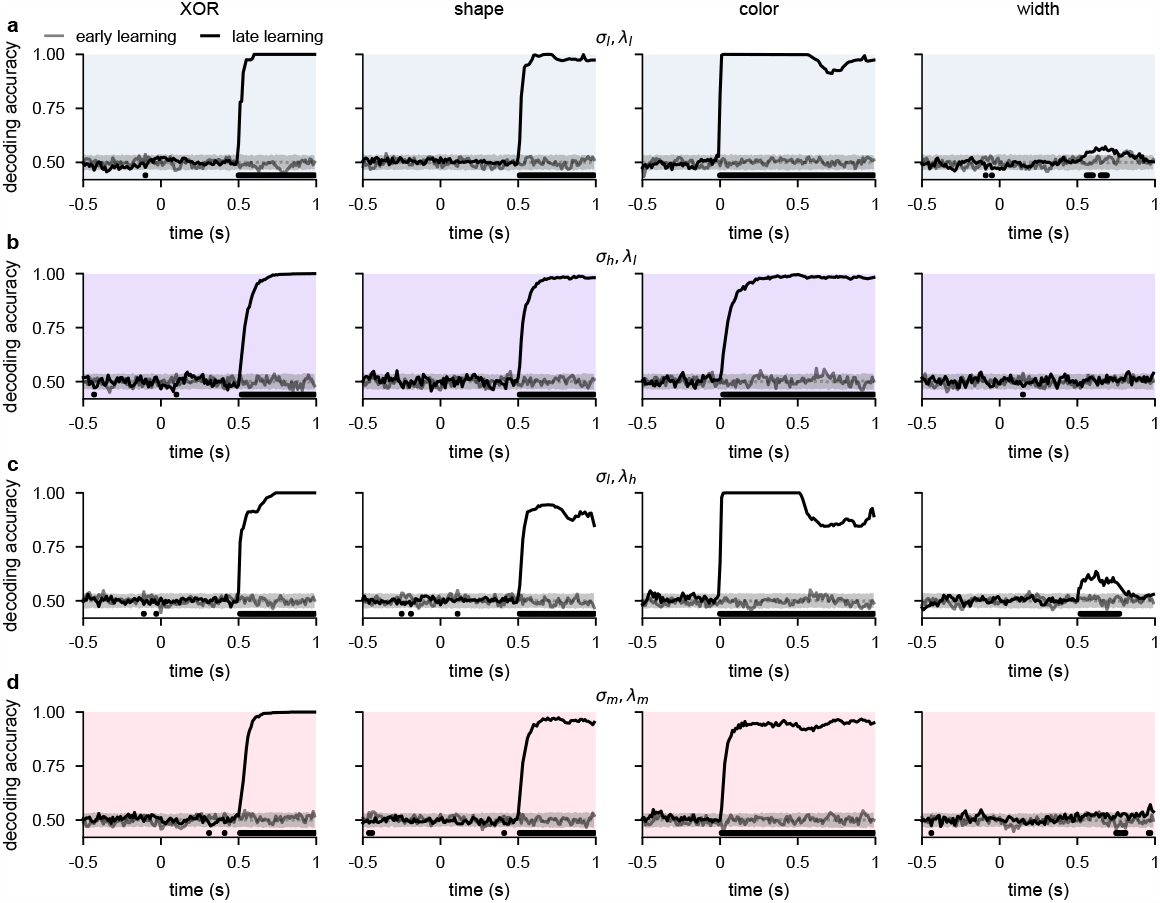
Temporal decoding for networks with input weights initialized to 0 prior to optimization. **a**, Performance of a linear decoder trained at each time point to predict either XOR (far left), shape (middle left), color (middle right), or width (far right) from neural population activity from optimized recurrent neural networks trained in the *σ*_l_, *λ*_l_ regime with input weights initialized to 0 prior to optimization (cf. Fig. 3a,b; Methods 1.2.1). We show decoding results separately from the first half of sessions (gray, ‘early learning’) and the second half of sessions (black, ‘late learning’). Dotted gray lines and shading show mean *±* 2 s.d. of chance level decoding based on shuffling trial labels. Horizontal black bars show significant differences between early and late decoding using a two-sided cluster-based permutation test (Methods 1.4). **b**, Same as panel a, but for networks optimized in the *σ*_h_, *λ*_l_ regime. **c**, Same as panel a, but for networks optimized in the *σ*_l_, *λ*_h_ regime. **d**, Same as panel a, but for networks optimized in the *σ*_m_, *λ*_m_ regime.

**Supplementary Fig. 4.**
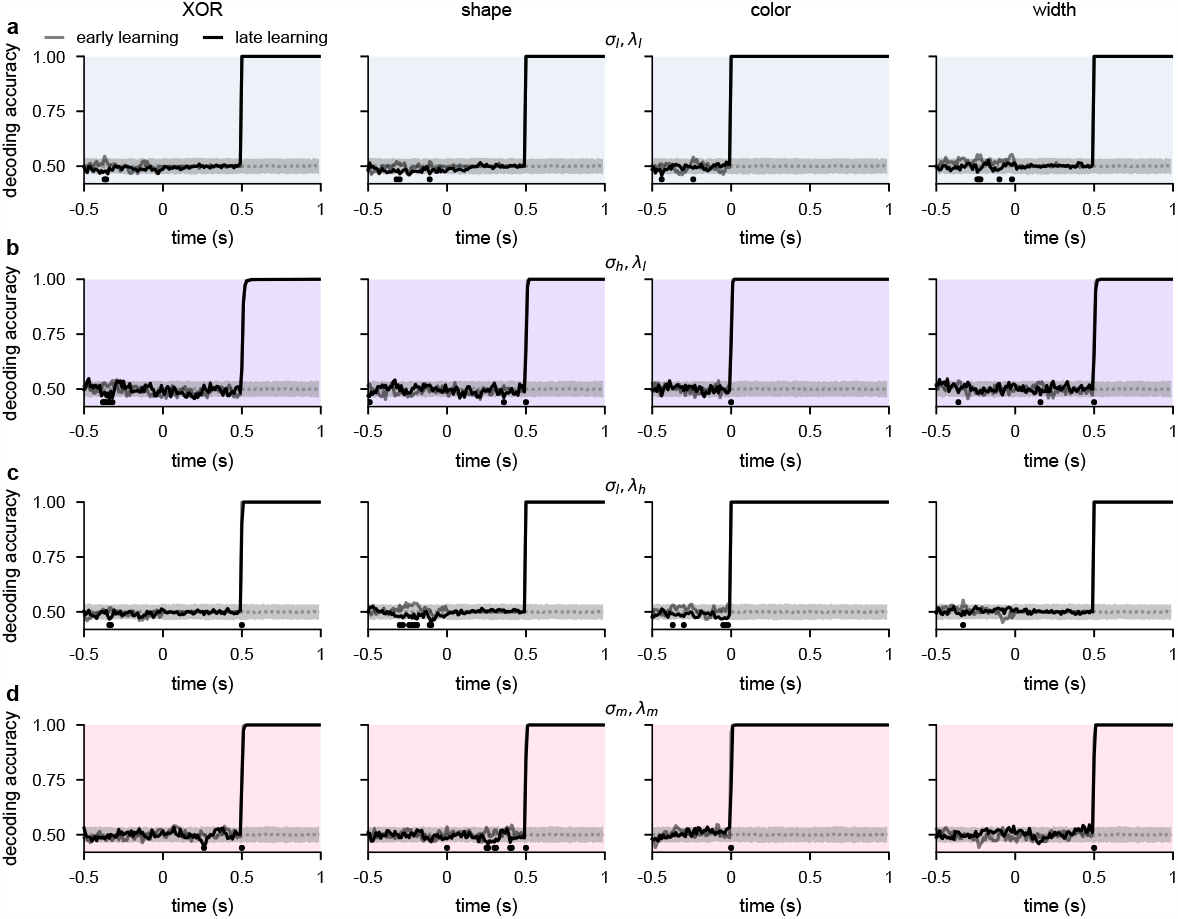
Temporal decoding for networks with input weights initialized to large random values prior to optimization. **a**, Performance of a linear decoder trained at each time point to predict either XOR (far left), shape (middle left), color (middle right), or width (far right) from neural population activity from optimized recurrent neural networks trained in the *σ*_l_, *λ*_l_ regime with input weights initialized to large random values prior to optimization (cf. Fig. 3a,b; Methods 1.2.1). We show decoding results separately from the first half of sessions (gray, ‘early learning’) and the second half of sessions (black, ‘late learning’). Dotted gray lines and shading show mean*±* 2 s.d. of chance level decoding based on shuffling trial labels. Horizontal black bars show significant differences between early and late decoding using a two-sided cluster-based permutation test (Methods 1.4). **b**, Same as panel a, but for networks optimized in the *σ*_h_, *λ*_l_ regime. **c**, Same as panel a, but for networks optimized in the *σ*_l_, *λ*_h_ regime. **d**, Same as panel a, but for networks optimized in the *σ*_m_, *λ*_m_ regime.

**Supplementary Fig. 5.**
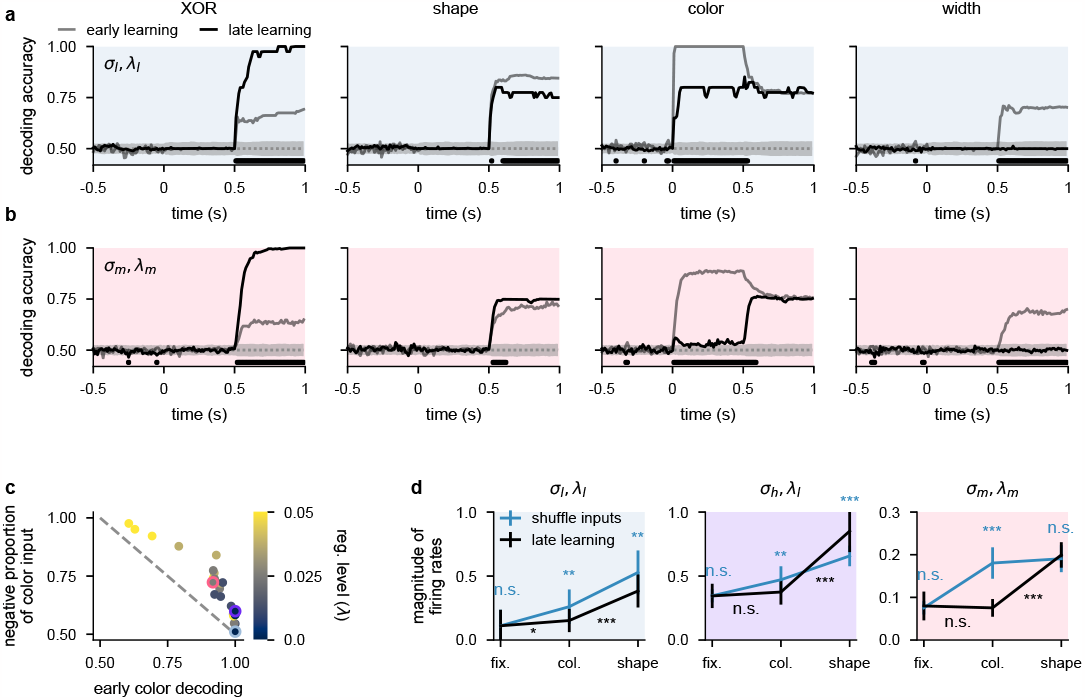
Temporal decoding for 2-neuron networks and supplemental analysis of suppression of dynamically irrelevant stimuli. **a**, Performance of a linear decoder trained at each time point to predict either XOR (far left), shape (middle left), color (middle right), or width (far right) from neural population activity from optimized 2-neuron networks (Fig. 5a–d) trained in the *σ*_l_, *λ*_l_ regime. Dotted gray lines and shading show mean *±* 2 s.d. of chance level decoding based on shuffling trial labels. Horizontal black bars show significant differences between early and late decoding using a two-sided cluster-based permutation test (Methods 1.4). **b**, Same as panel a but for 2-neuron networks trained in the *σ*_m_, *λ*_m_ regime. **c**, Mean (over 10 50-neuron networks) proportion of color inputs that have a negative sign for all noise and regularization strengths (colorbar) plotted against early color decoding (see Fig. 2c, middle). Pale blue, purple, and pink highlights correspond to the noise and regularization strengths shown in Fig. 2c. Gray dotted line shows the negative identity line shifted to intercept the vertical axis at 1. **d**, Momentary magnitude of firing rates 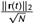; i.e. momentary metabolic cost, see Eq. 4 ) for 50-neuron networks in the *σ*_l_, *λ*_l_ regime (left, blue shading), *σ*_h_, *λ*_l_ regime (purple shading), and *σ*_m_, *λ*_m_ regime (right, pink shading). We show results after learning (black lines, ‘late learning’; repeated from Fig. 5f) and when shuffling color, shape, and width input weights (blue lines, ‘shuffle inputs’; Methods 1.3.3). Error bars show the mean*±* 2 s.d. over 10 different networks. P-values resulted from a two-sided Mann–Whitney U test (*, *p <* 0.05; **, *p <* 0.01; ***, *p <* 0.001; n.s., not significant; see Methods 1.4).

**Supplementary Fig. 6.**
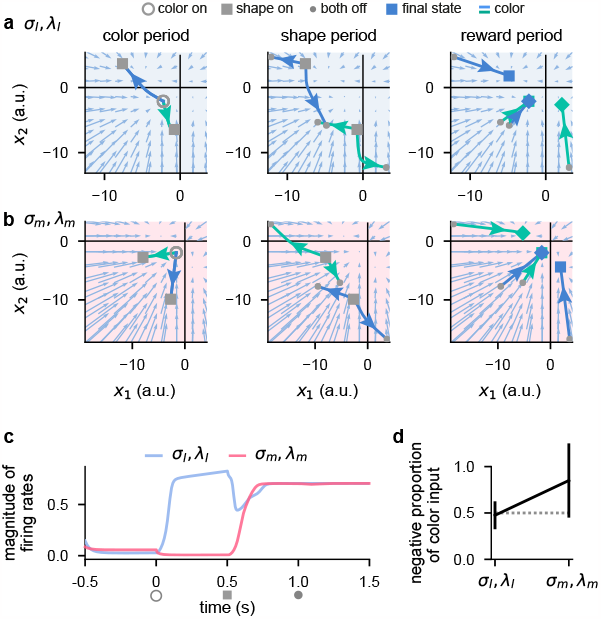
Activity-silent, sub-threshold dynamics lead to the suppression of dynamically irrelevant stimuli in networks with a sigmoid nonlinearity. **a**, Sub-threshold neural activity (**x**(*t*) in Eq. 1) in the full state space of an example 2-neuron network with a sigmoid (tanh) nonlinearity over the course of a trial (from color onset) trained in the *σ*_l_, *λ*_l_ regime (Methods 1.2). Pale blue arrows show flow field dynamics (direction and magnitude of movement in the state space as a function of the momentary state). Open gray circles indicate color onset, gray squares indicate shape onset, filled gray circles indicate offset of both stimuli, and colored squares and diamonds indicate the end of the trial at 1.5 s. We plot activity separately for the 3 periods of the task (color period, left; shape period, middle; reward period, right). We plot dynamics without noise for visual clarity. **b**, Same as panel a but for a network trained in the *σ*_m_, *λ*_m_ regime. **c**, Momentary magnitude of firing rates 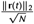 ; i.e. momentary metabolic cost, see Eq. 4 ) for the 2-neuron networks from panels a (blue line) and b (pink line). **d**, Mean*±* 2 s.d. (over 10 2-neuron tanh networks) proportion of color inputs that have a negative sign for the two noise and regularization regimes shown in panels a–c. Gray dotted line shows chance level proportion of negative color input.

