## Supplementary appendix for "Effects of noise and metabolic cost on cortical task representations"

**A1 Mathematical analysis of relevant and irrelevant stimulus coding in a linear network** . . . . . . . . . . **2**

A1.1 Problem definition 2

A1.2 The optimal linear decoder 2

A1.3 The performance of the optimal linear decoder 3

A1.4 Metabolic cost 4

A1.5 Qualitative predictions about optimal parameters 4

A1.6 Curvature of the loss function around the optimum 6

**References** . . . . . . . . . . . . . . . . . . . . . . . . . . . . . . . . . . . . . . . . . . . . . . . . **9**

In this supplementary appendix, we derive the optimal linear decoder for a linear network that receives both relevant and irrelevant inputs (Supplement A1.1 and A1.2). We derive how both the performance of the optimal decoder (Supplement A1.3) and the metabolic cost of the network (Supplement A1.4) depend on noise and the strength of relevant and irrelevant inputs. We also derive how the optimal setting of the relevant and irrelevant inputs, when jointly optimizing for both performance and a metabolic cost, depend on noise and the strength of metabolic cost (Supplement A1.5). Finally, we also study how noise and the strength of metabolic cost affect the curvature of the loss function around the optimum (Supplement A1.6).

### A1 Mathematical analysis of relevant and irrelevant stimulus coding in a linear network

Even though in the main text we simulate and numerically analyze neural networks with nonlinear temporal dynamics that solve an XOR task, for analytical tractability, our mathematical analyses below are for a model in which responses to different inputs combine linearly. Our analysis is agnostic as to whether responses come about by simple, instantaneous feedforward or temporally extended recurrent interactions, and is only concerned with the phenomenological mapping between stimuli and the resulting response distributions. As such, we cannot distinguish between dynamically and statically irrelevant stimuli, and so we only include a single relevant and single (statically) irrelevant stimulus in our analysis. Nevertheless, as we show in the main text (Fig. 4), some of the key insights from our analysis of such simplified systems generalize to the original setting.

#### A1.1 Problem definition

We consider the responses, $\mathbf{r}$, of a neural population of $N$ neurons to a stimulus that has a 'relevant' feature, $c$, which we assume to be binary with values $c_1$ and $c_2$ (occurring with uniform frequency), and to a (scalar) irrelevant feature, $\epsilon$ that is continuous (for convenience). We further assume that the responses are determined by a linear interaction of the relevant and irrelevant features and also subject to unspecific neural noise:

$$\mathbf{r} = \boldsymbol{\mu} + \delta_{c,c_1}\frac{\Delta\mathbf{r}_{\text{rel}}}{2} - \delta_{c,c_2}\frac{\Delta\mathbf{r}_{\text{rel}}}{2} + \epsilon\frac{\Delta\mathbf{r}_{\text{irrel}}}{2} + \boldsymbol{\eta} \tag{S1}$$

where $\boldsymbol{\mu}$ is the grand average response, averaging across both the relevant and the irrelevant feature, $\Delta\mathbf{r}_{\text{rel}}$ is the relevant tuning of the population, $\Delta\mathbf{r}_{\text{irrel}}$ is the irrelevant tuning, $\epsilon$ is (without loss of generality) zero mean and unit variance variability in the irrelevant feature, and $\boldsymbol{\eta} \sim \mathcal{N}\left(0, \sigma^2\,\mathbf{I}\right)$ is other neural noise.

In this note, we study how this neural population can optimise the decodability of $c$ while also balancing metabolic cost (defined below). Specifically, we will regard $\boldsymbol{\mu}$ and $\sigma^2$ as givens (constraints[1]) and ask how the population should "choose" $\Delta\mathbf{r}_{\text{rel}}$ and $\Delta\mathbf{r}_{\text{irrel}}$ (or some summary statistics thereof, see below) for this trade-off.

#### A1.2 The optimal linear decoder

We first study in general how information can be decoded from the population. The statistically optimal decoder, decoding $c$ from $\mathbf{r}$, is the maximum likelihood decoder (note that we have assumed $c_1$ and $c_2$ to occur with equal probabilities, see above). To make the derivations tractable, from here on we assume that $\epsilon$ is specifically Gaussian distributed, i.e. $\epsilon \sim \mathcal{N}(0,1)$, unless otherwise noted. In this case, the distribution of responses conditioned on the relevant stimulus feature becomes (equivariate) Gaussian, with some effective noise covariance $\boldsymbol{\Sigma}$ (to be determined later, Eq. S13):

$$\mathbf{r}|c_1 \sim \mathcal{N}\left(\boldsymbol{\mu} + \frac{\Delta\mathbf{r}_{\text{rel}}}{2}, \boldsymbol{\Sigma}\right) \tag{S2}$$

$$\mathbf{r}|c_2 \sim \mathcal{N}\left(\boldsymbol{\mu} - \frac{\Delta\mathbf{r}_{\text{rel}}}{2}, \boldsymbol{\Sigma}\right) \tag{S3}$$

Thus, the log odds ratio is

$$z = \ln\frac{\mathcal{P}\left(\mathbf{r}|c_1\right)}{\mathcal{P}\left(\mathbf{r}|c_2\right)} = \Delta\mathbf{r}_{\text{rel}}^{\top}\,\boldsymbol{\Sigma}^{-1}\,\mathbf{r} - \Delta\mathbf{r}_{\text{rel}}^{\top}\,\boldsymbol{\Sigma}^{-1}\,\boldsymbol{\mu} \tag{S4}$$

and the optimal decoder responds "1" if

$$z > 0 \tag{S5}$$

Thus, in this case, the optimal decoder is linear in $\mathbf{r}$, or conversely, a linear decoder (with the correct coefficients) is optimal.

[1] One could also consider $\boldsymbol{\mu}$ as an optimizable parameter, and its optimal value would simply be $\mathbf{0}$ (see footnote 3).

#### A1.3 The performance of the optimal linear decoder

We now turn to the question of how the parameters of the neural population affect its decodability, i.e. the performance of the optimal linear decoder. (We still assume that $\epsilon$, and thus $\mathbf{r}|c_{1/2}$ is Gaussian distributed.)

As we saw, the optimal decoder can be described by a simple thresholding of the log odds (Eq. S5). The log odds, $z$, itself is also Gaussian distributed conditioned on the relevant feature, because $\mathbf{r}$ is Gaussian distributed (Eqs. S2 and S3) and $z$ is a linear function of it (Eq. S4):

$$z|c_1 \sim \mathcal{N}\left(\mu_z, \sigma_z^2\right) \tag{S6}$$

and, due to symmetry,

$$z|c_2 \sim \mathcal{N}\left(-\mu_z, \sigma_z^2\right) \tag{S7}$$

with

$$\mu_z = \mathbb{E}\left[z|c_1\right] = -\mathbb{E}\left[z|c_2\right] = \frac{1}{2}\,\Delta\mathbf{r}_{\mathrm{rel}}^\top\,\mathbf{\Sigma}^{-1}\,\Delta\mathbf{r}_{\mathrm{rel}} \tag{S8}$$

and

$$\sigma_z^2 = \mathbb{V}\left[z|c_1\right] = \mathbb{V}\left[z|c_2\right] = \Delta\mathbf{r}_{\mathrm{rel}}^\top\,\mathbf{\Sigma}^{-1}\,\Delta\mathbf{r}_{\mathrm{rel}} = 2\,\mu_z \tag{S9}$$

The probability of correct decoding for $c_1$ (and by symmetry, also for $c_2$, and thus also after averaging over $c$) is given by

$$\Pi = \mathcal{P}\left(\text{respond “1”}|c_1\right) = \mathcal{P}\left(\text{respond “2”}|c_2\right) \tag{S10}$$

$$= \int_0^\infty \mathcal{N}\left(z;\mu_z,\sigma_z^2\right)\,\mathrm{d}z = \Psi\left(\frac{\mu_z}{\sigma_z}\right) = \Psi\left(\sqrt{\frac{\mu_z}{2}}\right) \tag{S11}$$

Therefore, the performance of the optimal linear decoder scales monotonically with $\mu_z$. To gain further insight into what this means in our particular setting, let us now express the effective noise covariance matrix $\mathbf{\Sigma}$ with the parameters of the problem definition:

$$\mathbf{\Sigma} = \mathbb{C}\left[\mathbf{r}|c_1\right] = \mathbb{C}\left[\mathbf{r}|c_2\right] \tag{S12}$$

$$= \sigma^2\,\mathbf{I} + \frac{\Delta\mathbf{r}_{\mathrm{irrel}}\,\Delta\mathbf{r}_{\mathrm{irrel}}^\top}{4} \tag{S13}$$

and its inverse (expressed using the Sherman-Morrison formula):

$$\mathbf{\Sigma}^{-1} = \frac{1}{\sigma^2}\left[\mathbf{I} - \frac{\Delta\mathbf{r}_{\mathrm{irrel}}\,\Delta\mathbf{r}_{\mathrm{irrel}}^\top}{4\,\sigma^2 + \Delta\mathbf{r}_{\mathrm{irrel}}^\top\,\Delta\mathbf{r}_{\mathrm{irrel}}}\right] \tag{S14}$$

Substituting Eq. S14 into the formula for $\mu_z$ (Eq. S8), we obtain:

$$\mu_z = \frac{1}{2\,\sigma^2}\,\Delta\mathbf{r}_{\mathrm{rel}}^\top\,\Delta\mathbf{r}_{\mathrm{rel}} - \frac{1}{2\,\sigma^2}\,\frac{\left(\Delta\mathbf{r}_{\mathrm{rel}}^\top\,\Delta\mathbf{r}_{\mathrm{irrel}}\right)\,\left(\Delta\mathbf{r}_{\mathrm{irrel}}^\top\,\Delta\mathbf{r}_{\mathrm{rel}}\right)}{4\,\sigma^2 + \Delta\mathbf{r}_{\mathrm{irrel}}^\top\,\Delta\mathbf{r}_{\mathrm{irrel}}} \tag{S15}$$

$$= \frac{1}{2\,\sigma^2}\,\left\|\Delta\mathbf{r}_{\mathrm{rel}}\right\|^2\left[1 - \frac{\left\|\Delta\mathbf{r}_{\mathrm{irrel}}\right\|^2}{4\,\sigma^2 + \left\|\Delta\mathbf{r}_{\mathrm{irrel}}\right\|^2}\left(\frac{\Delta\mathbf{r}_{\mathrm{rel}}^\top\,\Delta\mathbf{r}_{\mathrm{irrel}}}{\left\|\Delta\mathbf{r}_{\mathrm{rel}}\right\|\,\left\|\Delta\mathbf{r}_{\mathrm{irrel}}\right\|}\right)^2\right] \tag{S16}$$

Thus, $\mu_z$ can be simply expressed as

$$\mu_z = \frac{1}{2\,\sigma^2}\,\gamma^2\left(1 - \frac{\alpha^2}{4\,\sigma^2 + \alpha^2}\,\beta^2\right) \tag{S17}$$

where

$$\gamma = \left\|\Delta\mathbf{r}_{\mathrm{rel}}\right\| \text{ is the magnitude of tuning to the relevant feature} \tag{S18}$$

$$\alpha = \left\|\Delta\mathbf{r}_{\mathrm{irrel}}\right\| \text{ is the magnitude of tuning to the irrelevant feature} \tag{S19}$$

$$\text{and } \beta = \frac{\Delta\mathbf{r}_{\mathrm{rel}}^\top\,\Delta\mathbf{r}_{\mathrm{irrel}}}{\gamma\,\alpha} \text{ is the overlap of irrelevant with relevant tuning} \tag{S20}$$

which reveals that the effects of noise, relevant tuning, and irrelevant tuning factorise (corresponding to the first, second, and third terms, respectively), and that — intuitively — $\mu_z$ and thus performance increases with $\gamma$ and decreases with $\alpha$, $\beta$, and $\sigma^2$ (where we always consider the latter a constraint and thus fixed, see the problem definition).

Note that in the small noise limit, $\sigma^2 \ll \alpha^2$:

$$\mu_z = \frac{1}{2\,\sigma^2}\,\gamma^2\left(1-\beta^2\right) \tag{S21}$$

showing that performance in this case can only be increased by decreasing $\beta$ (or, trivially, by increasing $\gamma$), but not by decreasing $\alpha$. In contrast, when the small noise limit does not hold, the original Eq. S17 applies, and so performance can be increased by decreasing either $\alpha$ or $\beta$ (or, again, by increasing $\gamma$).

### A1.4 Metabolic cost

Following previous work [1–6], we define the metabolic cost to be the average sum of squared neural responses (where the averaging is over realizations of the relevant and irrelevant features as well as neural noise):

$$\omega^2 = \mathbb{E}\left[\|\mathbf{r}\|^2\right] \tag{S22}$$

which, by making the averaging over relevant features explicit, can be rewritten as

$$\omega^2 = \frac{\mathbb{E}\left[\|\mathbf{r}\|^2 \,|c_1\right] + \mathbb{E}\left[\|\mathbf{r}\|^2 \,|c_2\right]}{2} \tag{S23}$$

where the metabolic cost for $c_1$ is

$$\mathbb{E}\left[\|\mathbf{r}\|^2 \,|c_1\right] = \mathbb{E}\left[\mathbf{r}|c_1\right]^\top \mathbb{E}\left[\mathbf{r}|c_1\right] + \mathrm{Tr}\left(\mathbb{C}\left[\mathbf{r}|c_1\right]\right) \tag{S24}$$

$$= \left(\boldsymbol{\mu} + \frac{\Delta\mathbf{r}_{\mathrm{rel}}}{2}\right)^\top \left(\boldsymbol{\mu} + \frac{\Delta\mathbf{r}_{\mathrm{rel}}}{2}\right) + \mathrm{Tr}(\boldsymbol{\Sigma}) \tag{S25}$$

$$= \|\boldsymbol{\mu}\|^2 + \frac{\gamma^2}{4} + \boldsymbol{\mu}^\top \Delta\mathbf{r}_{\mathrm{rel}} + \frac{\alpha^2}{4} + N\sigma^2 \tag{S26}$$

and, again by symmetry,

$$\mathbb{E}\left[\|\mathbf{r}\|^2 \,|c_2\right] = \|\boldsymbol{\mu}\|^2 + \frac{\gamma^2}{4} - \boldsymbol{\mu}^\top \Delta\mathbf{r}_{\mathrm{rel}} + \frac{\alpha^2}{4} + N\sigma^2 \tag{S27}$$

and so, by substituting Eqs. S26 and S27 into Eq. S23, we get:

$$\omega^2 = \|\boldsymbol{\mu}\|^2 + N\sigma^2 + \frac{\gamma^2}{4} + \frac{\alpha^2}{4} \tag{S28}$$

This result is also interesting, because it shows that the metabolic cost factorises into four additive terms, each of which scales monotonically with a separate parameter: the average magnitude of responses, neural noise, the magnitude of relevant tuning, and the magnitude of irrelevant coding. It is also interesting to note what the metabolic cost does *not* depend on: the overlap between relevant and irrelevant tuning, $\beta$.

The first two terms in Eq. S28 are assumed to be fixed (see problem definition, Supplement A1.1), so we will not consider them further. The last two terms in Eq. S28 increase with $\gamma$ and $\alpha$, respectively. As we want the metabolic cost to be small, this prefers small $\gamma$ and $\alpha$.

Note that, unlike for deriving the optimal linear decoder (Supplement A1.2) and its performance (Supplement A1.3), for which we needed to assume that $\epsilon$ is normally distributed, no such assumption needed to be made for computing the metabolic cost (Eq. S28). Thus, for the metabolic cost, the same result is obtained for example when the irrelevant feature is binary, such that $\epsilon = \pm 1$ with equal probability (without loss of generality)[2], i.e. when variability in the irrelevant feature simply changes the responses by $\pm\Delta\mathbf{r}_{\mathrm{irrel}}$.

### A1.5 Qualitative predictions about optimal parameters

In general, the optimal setting of optimizable parameters, $\Delta\mathbf{r}_{\mathrm{rel}}$ and $\Delta\mathbf{r}_{\mathrm{irrel}}$ (Supplement A1.1), depends on the overall objective function, which will usually be a sum of performance (which we want to be high) and (negative) metabolic cost

[2] Note that, in this case, it is still true that $\mathbb{E}[\epsilon] = 0$ and $\mathbb{V}[\epsilon] = 1$, which is all what we assumed in the derivation above (see Eq. S13).

(which we want to be low), with some suitable Lagrange multiplier, $\lambda$, controlling the trade-off between the two terms:

$$\mathcal{L}(\gamma, \alpha, \beta) = \underbrace{\Pi}_{\text{Eqs. S11 and S17}} - \lambda \underbrace{\omega^2}_{\text{Eq. S28}} \tag{S29}$$

We note that the objective function also depends on the constraint parameters, $\boldsymbol{\mu}$ and $\sigma$ (Supplement A1.1). The effect of $\boldsymbol{\mu}$ is trivial: it only affects the metabolic cost, not performance, and it does so as a simple additive term (Eq. S28), so it does not affect the optimal values of the other parameters.[3] The effect of the other constraint, $\sigma$ is more nuanced, so we will separately consider two different regimes for it: small noise ($0 \leq \sigma \leq \sigma_{\text{crit}}$) and large noise ($\sigma_{\text{crit}} < \sigma$) with the constant $\sigma_{\text{crit}}$ defined later.

We also note that both performance (Eq. S17) and metabolic cost (Eq. S28) only depend on the optimizable parameters, $\Delta\mathbf{r}_{\text{rel}}$ and $\Delta\mathbf{r}_{\text{irrel}}$, through their summary statistics, $\gamma$, $\alpha$, and (for performance) $\beta$ (Eqs. S18-S20). In general, the dependence on the latter two parameters is straightforward: performance decreases in both $\alpha$ and $\beta$ (so that the last term in Eq. S17 achieves its maximal possible value 1 when either parameter is 0), while the metabolic cost increases with $\alpha$. This means that their optimal values will be at 0. However, we recall that in the small noise limit, only decreasing $\beta$ can improve performance (Eq. S21), while otherwise, decreasing $\alpha$ can also make a contribution to it. At the same time, the metabolic cost only depends on $\alpha$ but not on $\beta$. Thus to summarize, in the small noise limit, there is 'pressure' on both $\beta$ (from performance) and $\alpha$ (from the metabolic cost) to be small. In other cases, the pressure on $\alpha$ comes from both performance and metabolic cost, while $\beta$ only matters for performance, so we expect $\alpha$ to be more aggressively minimized by optimization, and perhaps $\beta$ not so much (which we find to be the case for our optimized recurrent neural networks; Fig. 4f,g; see also Supplement A1.6).

The overall effect of $\gamma$ is slightly less trivial. Eq. S28 shows that the metabolic cost depends on it quadratically, i.e. it should be small. However, as we saw earlier (Eq. S17), decoding performance monotonically grows with it. Nevertheless, this dependence of decoding performance on $\gamma$ is nonlinear (through the standard normal c.d.f., Eq. S11), such that it is effectively linear in $\gamma$ for small values[4], but saturates for large values of $\gamma$. This means that for small values of $\gamma$ the linear performance will dominate over the quadratic metabolic cost, but eventually, for large values, the quadratic metabolic term is guaranteed to dominate over the saturating performance, thus effectively limiting the magnitude of relevant tuning to some finite value. Note, however, that this argument does not yet reveal what happens to $\gamma$ depending on the noise regime, and where the transition between these two regimes happens.

Fortunately, it is possible to analytically derive $\gamma_*$, the optimal value of $\gamma$, as a function of $\sigma$. For this, we assume that $\alpha$ and $\beta$ already take their optimal values (without loss of generality, as we are interested in jointly optimising all relevant parameters), such that the last term in Eq. S17 is simply 1 (see above). In this case, the terms in the overall objective function (Eq. S29) that depend on the two remaining parameters of interest ($\gamma$ and $\sigma$) are simply:

$$\mathcal{L} = \Psi\left(\frac{\gamma}{2\,\sigma}\right) - \frac{\lambda}{4}\gamma^2 - \lambda\,N\,\sigma^2 + \ldots \tag{S30}$$

The optimal $\gamma$ can be simply defined implicitly as a function of $\sigma$ as the solution to the following equation:

$$0 = \frac{\partial\mathcal{L}}{\partial\gamma}(\gamma_*, \sigma) \tag{S31}$$

We now substitute (the partial derivative of) Eq. S30 into Eq. S31 to obtain:

$$0 = \frac{1}{2\,\sigma}\mathcal{N}\left(\frac{\gamma_*}{2\,\sigma}\right) - \frac{\lambda}{2}\gamma_* \tag{S32}$$

Note that this yields a solution for any $\sigma, \lambda > 0$ since the line $\frac{\lambda}{2}\gamma_*$ intersects the pdf of the Normal distribution for some $\gamma_* \geq 0$. Re-arranging Eq. S32 gives us

$$\gamma_*\, e^{\frac{\gamma_*^2}{8\,\sigma^2}} = \frac{1}{\sqrt{2\,\pi}\,\sigma\,\lambda} \tag{S33}$$

This equation has a solution in terms of the Lambert W function (defined by its inverse as $W^{-1}(x) = x\,e^x$)

$$\gamma_* = 2\,\sigma\sqrt{W\left(\frac{1}{8\,\pi\,\lambda^2\,\sigma^4}\right)} \tag{S34}$$

---

[3] This remains true even if $\boldsymbol{\mu}$ is optimisable, in which case it is also obvious from Eq. S28 that its optimal value is $\mathbf{0}$.

[4]This is because $\Psi(x)$ is linear in the $x \to 0$ limit, and the argument of $\Psi(\cdot)$ for computing the performance is linear in the square root of $\mu_z$ (Eq. S11), which in turn is quadratic in $\gamma$ (Eq. S17), so the argument of $\Psi(\cdot)$ is linear in $\gamma$.

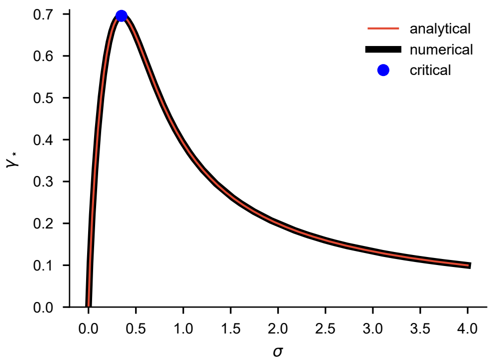


**Figure A1** |Plot of $\gamma_*$ as a function of $\sigma$ obtained by numerically optimising Eq. S30 (black), or using the analytical expression in Eq. S34 (red). Blue dot shows (analytically computed) critical values where $\gamma_*$ has a maximum (Eqs. S36 and S37). We used $\lambda = 1$ for these results.

where we only take the positive solution since $\gamma$ can only be positive.

We can find the maximum of this function by taking the derivative of Eq. S34 and setting it equal to 0. This gives:

$$W\left(\frac{1}{8\,\pi\,\lambda^2\,\sigma_{\text{crit}}^4}\right) = 1 \tag{S35}$$

Using the fact that $W(e) = 1$, we obtain

$$\sigma_{\text{crit}} = \frac{1}{(8\,\pi\,\lambda^2\,e)^{1/4}} \tag{S36}$$

Substituting this into Eq. S34 gives:

$$\gamma_{\text{crit}} = 2\,\sigma_{\text{crit}} \tag{S37}$$

These results are shown in Fig. A1, together with a numerical confirmation (see also Fig. 4d).

The qualitative dependence of $\gamma_*$ on $\sigma$ is straightforward to interpret and intuitive. In the *small* noise regime, $0 \leq \sigma \leq \sigma_{\text{crit}}$, $\gamma$ needs to grow with $\sigma$ so that the separation of the response distributions conditioned on $c_1$ and $c_2$ remains large enough to guarantee high performance. This growth starts at zero because for zero noise any infinitesimal separation between the mean responses for $c = c_1$ and $c_2$ makes for perfect performance. Thus, $\gamma$ remains small, and so — as we saw above — the performance term in the objective function dominates over the metabolic cost term. This means that the value of $\gamma$ that optimizes the full objective function can be understood from just this performance-based perspective. However, in the *large* noise regime, $\sigma_{\text{crit}} < \sigma$, $\gamma$ would need to be so large to guarantee high performance that it would reach a regime in which — as we saw above — the metabolic cost dominates. Thus the optimal $\gamma$ is increasingly influenced by the metabolic cost, and thus decreases with $\sigma$.

Finally, we derive classification performance with optimised parameters as a function of $\sigma$ by substituting Eq. S34 into Eqs. S11 and S17 and obtain

$$\Pi = \Psi\left(\sqrt{W\left(\frac{1}{8\,\pi\,\lambda^2\,\sigma^4}\right)}\right) \tag{S38}$$

which shows the intuitive result that performance monotonically decreases with $\sigma$ and drops to chance level for $\sigma \to \infty$ (Fig. A2, red). This is because the argument of $W$ monotonically decreases with $\sigma$ from infinity to zero, while both $W(x)$ and $\sqrt{x}$ monotonically increase (without bounds) with $x$ from zero at $x = 0$, and finally $\Psi(x)$ also monotonically increases with $x$ but from $\frac{1}{2}$ at $x = 0$ and to an asymptotic bound of 1.

### A1.6 Curvature of the loss function around the optimum

The curvature of the loss landscape around the optimum is important for determining how tightly constrained the parameters will be in the presence of noisy gradient updates due to the finite batch sizes used for stochastic gradient descent.

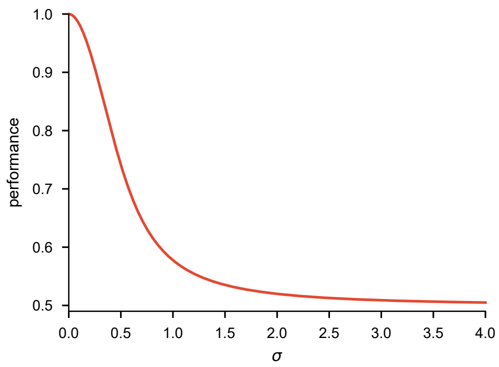


**Figure A2** |Plot of $\mathcal{P}$(correct) as a function of $\sigma$ when using $\gamma = \gamma_*$ that is optimised as in Fig. A1 (Eq. S38). We used $\lambda = 1$ for these results. All other parameters were optimised in all cases.

Before we proceed, we first repeat here the main equations regarding the loss $\mathcal{L}$:

$$\mathcal{L} = \Pi - \lambda\,\omega^2 \tag{S39}$$

$$\Pi = \Psi\left(\sqrt{\frac{\mu_z}{2}}\right) \tag{S40}$$

$$\mu_z = \frac{1}{2\,\sigma^2}\,\gamma^2\,\left(1 - \frac{\alpha^2}{4\,\sigma^2 + \alpha^2}\,\beta^2\right) \tag{S41}$$

$$\omega^2 = \|\boldsymbol{\mu}\|^2 + N\,\sigma^2 + \frac{\gamma^2}{4} + \frac{\alpha^2}{4} \tag{S42}$$

#### A1.6.1 Curvature of the loss function with respect to $\alpha$

In this section, we are interested in the first non-zero term of the Taylor expansion of the loss function $\mathcal{L}$ with respect to $\alpha$ — the magnitude of the irrelevant input — around the optimum at $\alpha = \beta = 0$. It turns out that the first non-zero term results from the second derivative of the metabolic cost term in the loss function since the other terms always contain at least an $\alpha$ or $\beta$ (which are 0 at the optimum). We therefore find that

$$\frac{\partial^2\mathcal{L}}{\partial\alpha^2}\Big|_{\alpha=0,\beta=0} = \frac{\partial^2\Pi}{\partial\alpha^2}\Big|_{\alpha=0,\beta=0} - \lambda\,\frac{\partial^2\omega^2}{\partial\alpha^2}\Big|_{\alpha=0,\beta=0} \tag{S43}$$

$$= 0 - \lambda\,\frac{\partial^2}{\partial\alpha^2}\left[\|\boldsymbol{\mu}\|^2 + N\,\sigma^2 + \frac{\gamma^2}{4} + \frac{\alpha^2}{4}\right] \tag{S44}$$

$$= -\frac{1}{2}\lambda. \tag{S45}$$

From this result, we see that the magnitude of the curvature of the loss decreases as a function of $\lambda$. In other words, the loss landscape as a function of the irrelevant input $\alpha$ becomes steeper with increasing regularization. This is why we observed larger values of $\alpha = \|\Delta\mathbf{r}_{\text{irrel}}\|$ (Eq. S18) in Fig. 4f with decreasing $\lambda$.

#### A1.6.2 Curvature of the loss function with respect to $\beta$

We now turn our attention to the first non-zero term of the Taylor expansion of the loss function $\mathcal{L}$ with respect to $\beta$ — the overlap of irrelevant with relevant tuning — around the optimum at $\alpha = \beta = 0$. It turns out that the first non-zero term results from a fourth derivative of the performance term $\Pi$ with respect to both $\alpha$ and $\beta$:

$$\frac{\partial^4\mathcal{L}}{\partial\alpha^2\,\beta^2}\Big|_{\alpha=0,\beta=0} = \frac{\partial^4\Pi}{\partial\alpha^2\,\beta^2}\Big|_{\alpha=0,\beta=0} - \lambda\,\frac{\partial^4\omega^2}{\partial\alpha^2\,\beta^2}\Big|_{\alpha=0,\beta=0} \tag{S46}$$

$$= \frac{\partial^4\Pi}{\partial\alpha^2\,\beta^2}\Big|_{\alpha=0,\beta=0} - \lambda\,\frac{\partial^4}{\partial\alpha^2\,\beta^2}\left[\|\boldsymbol{\mu}\|^2 + N\,\sigma^2 + \frac{\gamma^2}{4} + \frac{\alpha^2}{4}\right]\Big|_{\alpha=0,\beta=0} \tag{S47}$$

$$= \frac{\partial^4\Pi}{\partial\alpha^2\,\beta^2}\Big|_{\alpha=0,\beta=0} - 0. \tag{S48}$$

$$= \frac{\partial^4\Pi}{\partial\alpha^2\,\beta^2}\Big|_{\alpha=0,\beta=0} \tag{S49}$$

To evaluate this term we note that:

$$\frac{\partial^2 \Pi}{\partial \beta^2} = \frac{\partial^2 \Pi}{\partial \mu_z^2}\frac{\partial \mu_z}{\partial \beta} + \frac{\partial \Pi}{\partial \mu_z}\frac{\partial^2 \mu_z}{\partial \beta^2} \tag{S50}$$

$$= \frac{\partial^2 \Pi}{\partial \mu_z^2}\left(-\frac{\gamma^2}{2\sigma^2}\frac{2\,\alpha^2\,\beta}{4\,\sigma^2 + \alpha^2}\right) + \frac{\partial \Pi}{\partial \mu_z}\left(-\frac{\gamma^2}{2\sigma^2}\frac{2\,\alpha^2}{4\,\sigma^2 + \alpha^2}\right) \tag{S51}$$

Therefore

$$\tag{S52}$$

$$\frac{\partial^4 \Pi}{\partial \alpha^2\,\beta^2}\Big|_{\alpha=0,\beta=0} = \frac{\gamma^2}{2\sigma^2}\frac{\partial^2}{\partial \alpha^2}\left(\frac{\partial^2 \Pi}{\partial \mu_z^2}\left(-\frac{2\,\alpha^2\,\beta}{4\,\sigma^2+\alpha^2}\right) + \frac{\partial \Pi}{\partial \mu_z}\left(-\frac{2\,\alpha^2}{4\,\sigma^2+\alpha^2}\right)\right)\Big|_{\alpha=0,\beta=0} \tag{S53}$$

$$= \frac{\beta\gamma^2}{2\sigma^2}\frac{\partial^2}{\partial \alpha^2}\left(\frac{\partial^2 \Pi}{\partial \mu_z^2}\left(-\frac{2\,\alpha^2}{4\,\sigma^2+\alpha^2}\right)\right) + \frac{\gamma^2}{2\sigma^2}\frac{\partial^2}{\partial \alpha^2}\left(\frac{\partial \Pi}{\partial \mu_z}\left(-\frac{2\,\alpha^2}{4\,\sigma^2+\alpha^2}\right)\right)\Big|_{\alpha=0,\beta=0} \tag{S54}$$

$$= \frac{\gamma^2}{2\sigma^2}\frac{\partial^2}{\partial \alpha^2}\left(\frac{\partial \Pi}{\partial \mu_z}\left(-\frac{2\,\alpha^2}{4\,\sigma^2+\alpha^2}\right)\right)\Big|_{\alpha=0,\beta=0} \tag{S55}$$

Now, noting that $\frac{\partial}{\partial \alpha} = \frac{\partial}{\partial \mu_z}\frac{\partial \mu_z}{\partial \alpha}$, we obtain

$$\frac{\partial^2}{\partial \alpha^2}\left(\frac{\partial \Pi}{\partial \mu_z}\left(-\frac{2\,\alpha^2}{4\,\sigma^2+\alpha^2}\right)\right)\Big|_{\alpha=0,\beta=0} = -\frac{2\,\alpha^2}{4\,\sigma^2+\alpha^2}\left(\frac{\partial^2 \Pi}{\partial \mu_z^2}\frac{\partial^2 \mu_z}{\partial \alpha^2} + \frac{\partial^3 \Pi}{\partial \mu_z^3}\left(\frac{\partial^2 \mu_z}{\partial \alpha}\right)^2\right) + \frac{\partial \Pi}{\partial \mu_z}\frac{\partial^2}{\partial \alpha^2}\left(-\frac{2\,\alpha^2}{4\,\sigma^2+\alpha^2}\right)\Big|_{\alpha=0,\beta=0} \tag{S56}$$

$$= \frac{\partial \Pi}{\partial \mu_z}\frac{\partial^2}{\partial \alpha^2}\left(-\frac{2\,\alpha^2}{4\,\sigma^2+\alpha^2}\right)\Big|_{\alpha=0,\beta=0} \tag{S57}$$

$$= \frac{\partial \Pi}{\partial \mu_z}\frac{\partial}{\partial \alpha}\left(-\frac{4\alpha}{4\sigma^2+\alpha^2} + \frac{4\alpha^3}{(4\sigma^2+\alpha^2)^2}\right)\Big|_{\alpha=0,\beta=0} \tag{S58}$$

$$= \frac{\partial \Pi}{\partial \mu_z}\left(-\frac{4}{4\sigma^2+\alpha^2} + \frac{8\alpha}{(4\sigma^2+\alpha^2)^2} + \frac{12\alpha^2}{(4\sigma^2+\alpha^2)^2} - \frac{16\alpha^4}{(4\sigma^2+\alpha^2)^3}\right)\Big|_{\alpha=0,\beta=0} \tag{S59}$$

$$= -\frac{4}{4\sigma^2}\frac{\partial \Pi}{\partial \mu_z}\Big|_{\alpha=0,\beta=0} \tag{S60}$$

$$= -\frac{4}{4\sigma^2}\frac{1}{4}\sqrt{\frac{2}{\mu_z}}\mathcal{N}\left(\sqrt{\frac{\mu_z}{2}}\right)\Big|_{\alpha=0,\beta=0} \tag{S61}$$

$$= -\frac{1}{4\sigma^2}\frac{2\sigma}{\gamma}\mathcal{N}\left(\frac{\gamma}{2\sigma}\right) \tag{S62}$$

$$= -\frac{1}{2\sigma\gamma}\mathcal{N}\left(\frac{\gamma}{2\sigma}\right) \tag{S63}$$

Therefore, from Eq. S55, we obtain

$$\frac{\partial^4 \mathcal{L}}{\partial \alpha^2\,\beta^2}\Big|_{\alpha=0,\beta=0} = -\frac{\gamma}{4\sigma^3}\mathcal{N}\left(\frac{\gamma}{2\sigma}\right) \tag{S64}$$

We therefore find that the magnitude of this term becomes smaller with increasing noise. This is why we observed larger values of $\beta = \frac{\Delta \mathbf{r}_{\text{rel}}^\top \Delta \mathbf{r}_{\text{irrel}}}{\|\Delta \mathbf{r}_{\text{rel}}\|\,\|\Delta \mathbf{r}_{\text{irrel}}\|}$ (Eq. S20) in Fig. 4g with increasing $\sigma$.
